# HIGH FREQUENCY ROOT DYNAMICS: SAMPLING AND INTERPRETATION USING REPLICATED ROBOTIC MINIRHIZOTRONS

**DOI:** 10.1101/2022.01.06.475082

**Authors:** Richard Nair, Martin Strube, Martin Hertel, Olaf Kolle, Victor Rolo, Mirco Migliavacca

## Abstract

Automating dynamic fine root data collection in the field is a longstanding challenge with multiple applications for co-interpretation and synthesis for ecosystem understanding. High frequency root data are only achievable with paired automated sampling and processing. However, automatic minirhizotron (root camera) instruments are still rare and data is often not collected in natural soils nor analysed at high temporal resolution. Instruments must also be affordable for replication and robust under variable natural conditions.

Here, we show a system built with off-the-shelf parts which samples at sub-daily resolution. We paired this with a neural network to analyse all images collected. We performed two mesocosm studies and two field trials alongside ancillary data collection (soil CO_2_ efflux, temperature and moisture content, and ‘PhenoCam’-derived above-ground dynamics. We produce robust daily time-series of root dynamics under all conditions. Temporal root changes were a stronger driver than absolute biomass on soil CO_2_ efflux in mesocosm. Proximal sensed above ground dynamics and belowground from minirhizotron data were not synchronised. Root properties extracted were sensitive to soil moisture and occasionally to time of day (potentially relating to soil moisture). This may only affect high frequency imagery and should be considered in interpreting such data.

**HIGHLIGHT:** Completely automatic root dynamics with method transferrable between field settings.

## INTRODUCTION

Plant phenology (seasonal patterns of recurrent events such as leaf growth and senescence) drives interannual variability of the terrestrial carbon (C) sink (Raupach *et al*., 2011). It responds to environmental conditions such as climate and weather (Richardson *et al*., 2013), can differ above and below ground (Adair *et al*., 2019), and is partially determined by life history (Steinaker *et al*., 2010). While overall growth is also driven by whole-plant resource budgets, root and shoot phenology are not always linked (Abramoff and Finzi, 2015). Above- and belowground-environments differ (e.g., Blume-Werry, Jansson, & Milbau, 2017) and plants temporally partition resource uptake and demand, resource assignment and activity of individual organs (Herrmann *et al*., 2016), which may cause this desynchrony.

Tools to remotely monitor ecosystem dynamics above-ground are well developed, for instance ‘PhenoCams’ (Luo *et al*., 2018; Richardson *et al*., 2018) or satellite-derived vegetation indexes (Wu et al., 2017). However, using above-ground measurements to proxy for below-ground activity is an unreliable assumption. Root phenology measurements are needed in natural field contexts and high frequency datasets are rare (Radville *et al*., 2016). This is because of the demanding, destructive, non-repeatable nature of traditional sampling. Data scarcity contributes to basic simulation of root dynamics by models, often relying on poorly calibrated root:shoot ratios, simple environmental triggers, or optimality concepts (De Kauwe *et al*., 2014; Walker *et al*., 2015). These are hard to validate without high frequency datasets in sites providing other data for model fitting.

Roots are also crucial for C cycling because they control decomposition, and are the main C source to soil organic matter (Dijkstra *et al*., 2021). Thus, misrepresentation of root dynamics affects prediction of ecosystem capacity to sequester carbon. Unfortunately, mesocosm experiments are prone to artefacts which disproportionately affect roots (Poorter *et al*., 2012) so the benefits of field experiments (Schindler, 1998) are particularly large for the below-ground. In natural field conditions, repeatable root observations are made with minirhizotrons (buried observatories and camera systems); many other advances in root phenotyping (e.g. Le Marié et al., 2014; Liu et al., 2021; Nagel et al., 2012) are only currently possible in more controlled conditions. Minirhizotron automation has been possible for 15 years (Allen *et al*., 2007) but still often use infrequent imaging and manual-driven analysis (e.g. Defrenne et al., 2021). Application of robotic minirhizotrons for monitoring below ground phenology of roots could offer many opportunities, but it has barely advanced beyond pioneering experiments using these systems (Vargas and Allen, 2008; Iversen *et al*., 2011; Allen and Kitajima, 2013, 2014).

Automated observations introduce a bottleneck: processing high frequency imagery to calculate proxies of root properties. A manual approach will inevitably not annotate images as fast as collection. Recently, convolutional neural networks (CNNs) showed promising results to identify roots in a variety of settings (Delory *et al*., 2016; Rahmanzadeh and Shojaedini, 2016; Vincent *et al*., 2016; Huo and Cheng, 2019; Bauer *et al*., 2022). In particular, field soil minirhizotron imagery has been analysed several times with good results (Wang *et al*., 2019; Smith *et al*., 2020; Gillert *et al*., 2021; Han *et al*., 2021; Peters *et al*., 2022; Bauer *et al*., 2022), but transferability between sites and out of agricultural soils is difficult to assess without widespread adoption in new settings. These have also never been applied to high frequency studies where variability between images (e.g. soil moisture, soil animals) may cause instability in timeseries data produced.

We built a robotic minirhizotron system (henceforth, RMR) with a per instrument parts budget of €2000, allowing replication. We paired this with an established CNN method *(Rootpainter*, Smith et al., 2020) previously used in low sampling frequency studies or soil cores (Han *et al*., 2021; Alonso-Crespo *et al*., 2022). We processed segmented images to extract basic root properties (root length, root length density) at high time resolution. Besides the collection step, there are three main challenges to using automated minirhizotrons: 1) usefulness of extracted properties to explain system functioning at fine timescales 2) comparability between different point sensors and 3) long term robustness in field operation. Automatic systems must work on schedule in adverse conditions, and image processing must be robust to non-opportune fixed sampling issues (e.g. condensation) which may be avoided in manual campaigns.

Here, we show O1) the RMR method is a reliable option to produce time series of basic architectural traits O2) consistent between instruments O3) under a variety of sometimes adverse conditions, and O4) interpretable in relation to system functioning. We did this through two mesocosm and two field experiments. In mesocosm: E1) using a single RMR in a greenhouse paired with proximal remote sensed above-ground dynamics and soil CO_2_ fluxes (O1,O4), and E2) eight replicated RMRs in a greenhouse paired with proximal remote sensed above-ground dynamics only (O1,O2). In the field: E3) for four months in the autumn in a Mediterranean tree-grass ecosystem (O1, O3, O4) and for E4) two months in winter/spring in a temperate grassland (O1, O3). Field experiments encountered heterogenous soil appearances, condensation, soil animals, root litter and other disturbances potentially affecting instrument operation, human annotation and consistency of CNN segmentation across a high frequency timeseries. These were also replicated but at a lower scale than E2 (O2).

## MATERIALS AND METHODS

### Minirhizotron System Design

We based our minirhizotron instrument (RMR, Fig. 1) on a movable camera design able to travel in two axes (along a minirhizotron tube, and rotationally around a minirhizotron tube). All components of the system were purchased unmodified, with the exception of a customised fish-eye lens to allow the camera to focus at (∼ 3 cm) within the observatory and the linear actuator which was built to our dimensions. The RMR uses an (internal) 10/9.6 (external/internal) x 100 cm observatory. This is comparable to the only other automatic minirhizotron systems used in published experiments (Iversen *et al*., 2011; Svane *et al*., 2019). It takes around 40 minutes to sample 112 separate images covering the entire tube. Timing of cycles is set by a timer switch and the instrument completes one sampling cycle whenever it is powered on. The light source is provided by a ring of LEDs which were angled away from the image subject to reduce reflectance. The instrument framework was made of black plastic and all exposed screw heads were painted black for the same purpose. Cable management was achieved by affixing cables to a rigid band only able to bend in the lateral direction. A power supply converter from 12V to 5V was used for the PC, camera and LED ring lights. One sampling cycle required around 10 Watt hours of power.

**Fig. 1.**
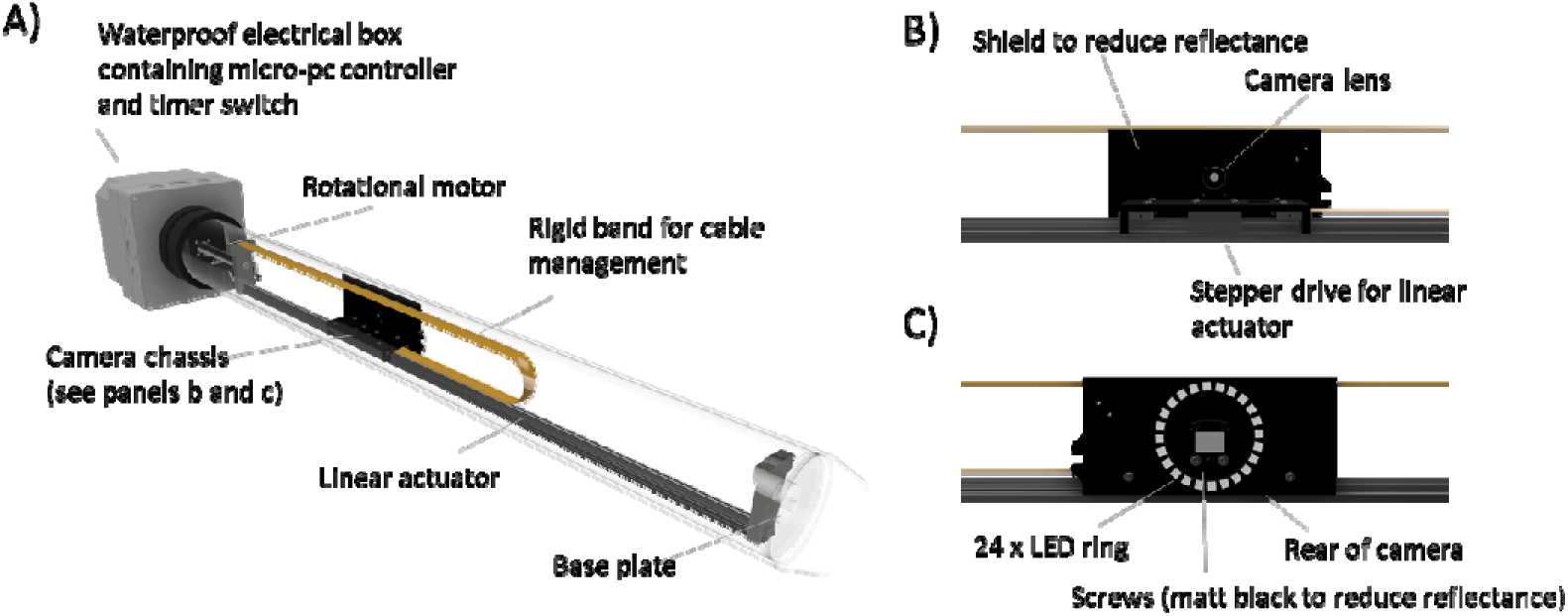
Conceptual diagram of the RMR. The full system (A) moves a camera chassis (B) longitudinally and rotationally to image soil in close contact with the tube. The camera chassis is designed to minimise reflectance by mounting the light source away from the focal area (C).

RMRs operate at any angle, we ran the RMR horizontally in the greenhouse (E1 and E2) or at an angle in the field (E3 and E4). We tested the RMR across a range of temperatures from -20°C to ∼ 35°C (see details later). Images (25 μm pixel^-1^, ∼1000 dpi. in our final system) are captured with a small overlap in .jpg format. These were saved onto removeable 128 GB SD cards; in order to obtain the images, the SD cards were swapped with blank alternatives when the instruments were not powered. A log file was amended at the end of each imaging cycle which could be checked much faster than individual images and easily accessed over WiFi. A single image was less than 1 Mb, therefore the RMR could sample around 1100 cycles without the SD card being changed. Further information about the design can be found in Supplementary Protocol S1 and a summary of the differences between experiments in Table 1.

**Table 1.** Summary of experiment objectives, camera hardware and duration.

| EXPERIMENT | OBJECTIVE | REVISIONS | N. INSTRUMENTS | DURATION |
| --- | --- | --- | --- | --- |
| E1 | Proof of Concept (Biological Interpretation) | Camera: IDS 1005XS-C<br>Mains Power | 1 | 3 months (greenhouse) |
| E2 | Cross-Instrument Consistency | Camera: DFK AFU050-L34<br>Mains Power | 8 | 2 months (greenhouse) |
| E3 | Field Trial | Camera: DFK AFU050-L34<br>Solar Power | 3 * | 4 months (October – January) |
| E4 | Field Trial (low temperature) | Camera: IDS 1007XS-C<br>Mains Power<br>GPS Clock<br>Magnetic 'home' positioning | 2 | 60 days (February – March) |

\* In E3, we ran eight instruments in the field, but had problems with long-term timekeeping due to low temperatures which limited this number to three concurrent instruments with good data. This issue was subsequently fixed for E4 but the instruments were split between two sites (second site not shown) to limit vulnerability to travel disruption.

### Greenhouse Experiments

We designed two greenhouse experiments: E1) with one RMR (128,880 individual images) and E2), with eight RMRs (216,000 individual images). In E1 we sampled every two hours and had additional ancillary instruments, in E2 we sampled every six hours without paired ancillary instruments. Each mesocosm (140 cm (L) x 40 cm (W) x 30 cm (H)) was filled with soil sieved to 0.5 cm (roots and stones removed), harvested from Meusebach, Germany (L. Eder pers. comm.). Soil C:N was 18:2. Soil covered the upmost surface of the RMR by 2 cm, so field of view spanned 2 to 12 cm below the soil surface.

In both experiments, a three species seed mix (80/15/5 % by mass *Anthroxanthum odoratum, Plantago lanceolata, Medicago scutellata*) was scattered evenly over the soil surface, kept moist to germinate. Water was provided at irregular but recorded intervals and volumes via a watering can with a distributor nozzle to mimic field rain events. We watered until an arbitrary date, then withheld water to move the system towards a state of drought-induced root senescence. This senescence period was longer in E2 (1 month) than E1 (2 weeks).

Each RMR observatory was located in a mesocosm unit (Fig. S2), extending internally 90 cm from one end. As here the RMR was horizontal, the first ∼ 10 cm of the length, usually partially above ground, was not sampled. Aside from the camera, which we changed because of a supply issue (resulting in worse image quality in E2), and minor modifications for cable management, the RMR design was the same. In E1 we used a ‘fresh’ observatory installed factory clean, but in E2 we re-used an experimental set up for manual measurements which we cleaned before starting. Thus, this latter experiment potentially had artefacts on the tube surface, which we discuss later. Both experiments were run in the greenhouse of the Max Planck Institute for Biogeochemistry, Jena, Germany.

All mesocosm units in both greenhouse experiments were included in the field of view of a standard ‘PhenoCam’ set up (Sonnentag *et al*., 2012; Richardson *et al*., 2018), modified for indoor use via a fish-eye lens. The camera settings were defined from the ‘PhenoCam’ protocol (Richardson *et al*., 2018). Images were collected between midday and 1pm local time. We defined a single ROI for each entire mesocosm unit, parallel to the long axis of the minirhizotron. Data was processed as in field-scale studies (e.g. Luo et al., 2018, 2020). During this time artificial lights were switched off and watering was never during this hour. Field of views (FOVs) during each experiment were stable and images were available from the entire period. From the images we obtained daily green chromatic coordinate (GCC), the ratio between digital numbers in the green channel divided by the sum of the digital numbers in red, green and blue channels, commonly used to represent canopy greenness in field studies. GCC has been found to outperform NDVI in forests when assessing vegetation cover and condition and supressing scene illumination variation (Nijland *et al*., 2014). Using greenness indexes assumes that healthy vegetation is greener than less healthy vegetation.

We also installed soil moisture and soil temperature probes (a combination of EC-5 soil moisture probes (Li-COR Biosciences, USA) and ML-3 soil temperature probes (Decagon Instruments, USA), and measured root biomass through six 4.5 cm x 13 cm ingrowth cores in each mesocosm installed at initiation. These were evenly spaced, at 5 cm from the wall of the unit and an equal distance from the central minirhizotron tube. These were retrieved and root biomass measured by sieving the soil to 2 mm and weighing the washed and dried (60°C, 3 days) roots. In E1, we sampled six un-replicated time points to minimize disturbance and because of the limited area of soil not occupied by the minirhizotron or the gas exchange measurement, plus the start (no roots), while in E2 we sampled the start (no roots), three un-replicated time points, and then one time point at the end of the experiment with 3 replicates per mesocosm.

Finally, in E1, we measured system gas exchange using a Li-8100A Infrared Gas Analyzer (IRGA), a Li-8100-104 opaque long-term chamber (Li-COR Biosciences, Lincoln, USA) every half-hour, processing these data via standard packages. We let vegetation grow within the chambers. These data were processed using the R package RespChamberProc; in general, the fit of all observations was very good (> 99% with an R^2^ of 0.99). Occasional, unplanned periods of power disruption in both experiments prevented data collection and image capture but did not affect the subsequent minirhizotron images.

### Field Trials

The third (E3, 68432 images, 203 cycles per instrument) and fourth trial (E4, 53760 images, 240 cycles per instrument) were in field settings. E3 was at Majadas de Tiétar,(Spain), a Mediterranean wood-pasture (El-Madany et al., 2018; Nair et al., 2019). We deployed eight RMRs as in E2 powered by per-instrument solar panels coupled with an external 12V battery and charge controller. We sampled twice daily from October 2019 until January 2020 (∼ 22800 images per instrument). Majadas de Tiétar is a Mediterranean ecosystem where the growing season lasts from autumn until late spring, but much undecomposed root litter remains after dry summer (Nair *et al*., 2019). The soil is an Abruptic Luvisol with a sandy upper layer and a thick clay layer starting at 20-40 cm. In E3, we encountered an unpredictable issue with the BIOS clock when night temperatures fell below 0°C, affecting accurate timekeeping (and hence reference time for data collected) at all subsequent sampling points. Therefore, we show summary data from two instruments where timekeeping was not disrupted across all four months and a third composite instrument made from two instruments with partial timeseries (i.e. only ‘true’ time referenced data used, with three instruments over 80 % of the timeseries, gapfilled via linear interpolation for periods missing data). We also use a unified meteorological dataset for a large-scale ecosystem experiment (‘MANIP’) at the site. Air temperature was the mean of three measurements made at 2 m height at three eddy covariance towers located within approximately 950 m of each other. Precipitation was likewise the mean of precipitation recorded at each tower. GCC used in this experiment was the mean of ‘pasture’ GCC captured from a ‘PhenoCam’ digital camera used for site comparison between the three towers (Luo et al., 2020) with a three-day average taken for a daily value.

E4 was conducted in plots of the Jena biodiversity Experiment (Roscher *et al*., 2004; Weisser *et al*., 2017) in Jena, Germany from 1 Feb to 04 April 2022 where mean air temp. was 4.5 °C, and minimum -7.5 °C. Here two instruments ran from mains power sampling four times daily. A single meteorological station was located in the centre of the experiment. Air temperature was measured at 2 m and rainfall from a single precipitation gauge. This site is a loamy Eutric Fluvisol in the floodplain of the river Saale. The RMR had minor modifications to ensure correct timekeeping at low temperatures and we replaced the camera used in E2 and E3 with another model. We further tested this robustness in a -20°C cold room (not shown). In E3 and E4 observatories were installed at 40°, E3 in May 2015 and E4 in August 2020. In this manuscript we show root properties averaged over the whole observatory depth, which in both E3 and E4 reached 45 cm underground. A summary of objectives and hardware differences between E1-E4 are in Table 1.

### Processing Minirhizotron Imagery for Root Traits

Images collected from all four experiments were very different and contained different artefacts (Fig. 2). We processed minirhizotron imagery in two steps: 1) segmentation of the images into binary maps using a CNN 2) post-processing of the binary images to extract root properties.

**Fig. 2.**
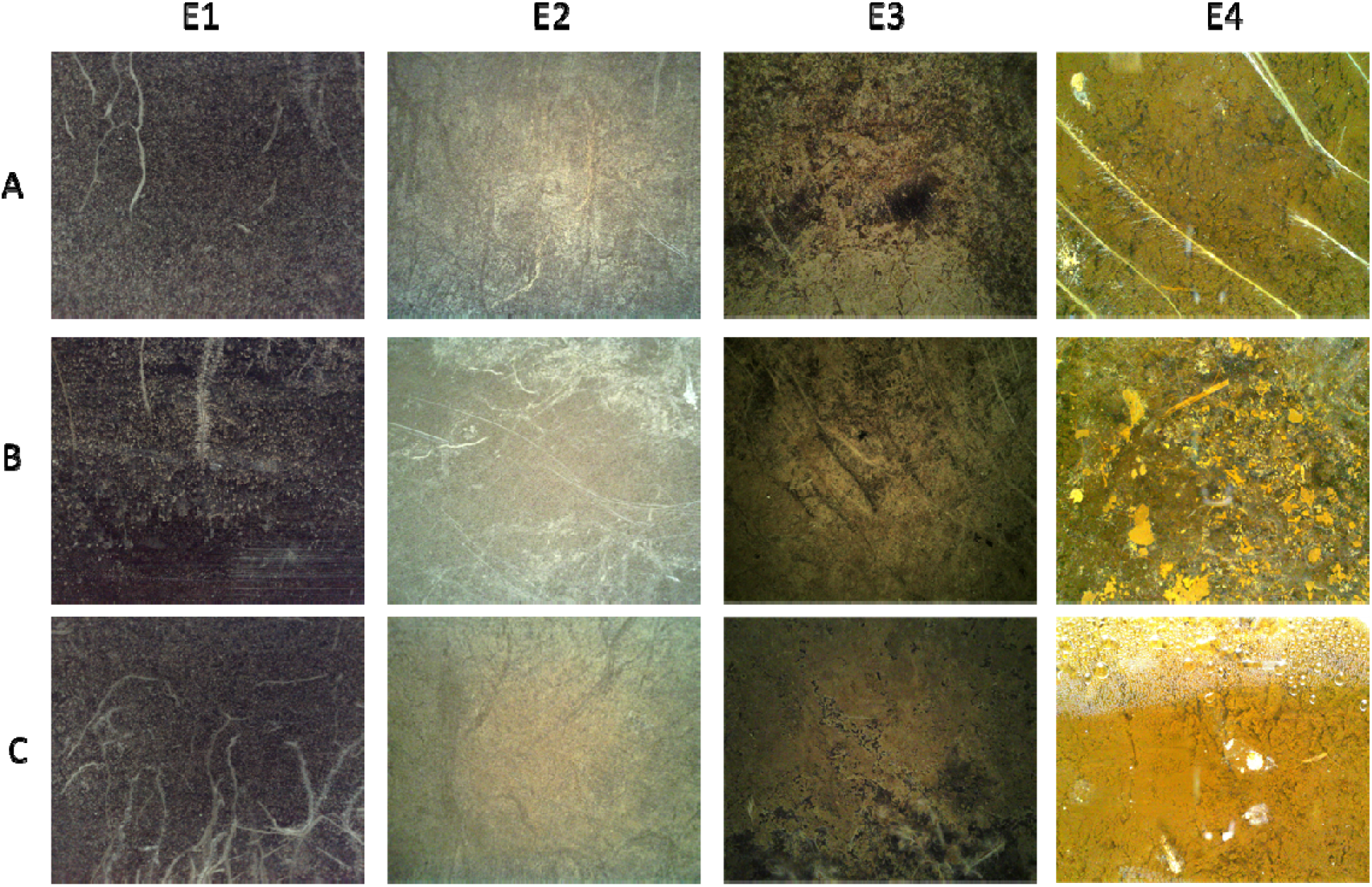
Example of 3 random images (A,B,C) from each of our experiments. Each image is originally 2292 × 1944 mm. The soil appearance and image quality were very different between experiments due to a combination of different image sensors and different image subjects. Notable examples of segmentation challenges are scratches and tube surface artefacts E1B,E2B), litter and remains of dead roots (E3), condensation and soil animals (E4C).

In step 1, we trained a CNN with a Graphical User Interface, allowing corrective annotation (*Rootpainter*, Smith et al., 2020). For this study, we hosted our own remote server, using local computer cluster GPU cores. We used a random model trained from scratch in each experiment. We aimed for separation of RMR image pixels into two classes, roots and soil. We started with a random model without any pre-annotated images, training on 308 (E1), 300 (E2), 400 (E3) or 350 (E4) complete 2292 × 1944 images. We stopped training when we (qualitatively) assessed the segmentation was not improving and had processed at least 300 images. Training data is a major issue for automatic minirhizotron studies which have only short intervals between images but may have sudden changes (e.g. changes in soil colour following rain). Random annotation is unlikely to capture these events and targeting these times introduces bias. Because of the camera change (Table 1), and potentially because of the legacy of previous roots around the observatory, images in E2 and E3 were also challenging to both human annotations and CNN segmentation due to low contrast, poor focus and faint marks on the tube surface. E3 also had high litter content, leading to ambiguity between roots and litter and rapid soil appearance changes as the ecosystem was released from summer drought. Thus, high throughput of training data via corrective annotation / active learning strategies (e.g. Beluch et al., 2018; Budd et al., 2021; Ren et al., 2021) offers a major advantage in such unpredictable tasks.

We then processed the segmented dataset to produce basic root traits. In this analysis we extracted root length and root surface area (RSA). To do this, we first applied a filter on the segmentation with a total area of less than 0.5 mm^2^, which we treated as noise. Because our minirhizotron images contain partial aspects of more than one plant’s root system, we defined root surface area as the total area remaining. We divided this by the area of the image for a value in percentage. For root length, we took the segmented image and post-procesed it through scripts in python 3. To calculate root length, we skeletonised all segmented images to one pixel wise, and calculated the total length from the number of pixels in the skeleton. We did not adjust for diagonal connections nor prune branches of the skeleton but otherwise this is similar to the simplest level of analysis using modern software for root imagery such as RhizovisionExplorer (Seethepalli *et al*., 2021). This software can be directly paired to the rootpainter CNN and has been successfully use to process imagery from agricultural soils with similar accuracy to manual annotation (Bauer *et al*., 2022) and can potentially extract many more features of interest. However, our datasets in natural ecosystems were both challenging to the CNN segmentation and for the human operator to choose appropriate settings for the software. We also faced a software constraint from the huge number of segmented images we had to process in one go restricting the use of GUI-based software. Hence, we used a simpler programmatic option to generate our timeseries where we had the advantage of frequent resampling to make up for individual inaccuracy. We further converted root length into root length density (RLD) by dividing the total root length by the total area of the image as we were only observing those portions of root which encountered the observatory and not whole roots. RLD is a common property in analyses relating to soil exploration and resource uptake from fine roots (Freschet *et al*., 2021), while RSA incorporates variation in diameter and can be assumed to correlate closer to observable biomass if density is stable.

To validate, we use 378/170/140/90 manually annotated images (E1/E2/E3/E4). Images were annotated by multiple users (E1,E2,E3) or the lead author (E4) with anonymous filenames in GIMP software (The GIMP Development Team, 2019). In E1 and E2 we tried to produce high quality annotation with up to an hour of annotation time per image. For E3 and E4 these were but a ‘fast-pass’ annotation (max. 10 min annotation time per image, often less) deemed suitable for high throughput timeseries. We compared at image level, i.e. we did not assess the fit of individual pixels. By working at the image level, we discarded the advantage the minirhizotrons have in tracking individual roots, because we were interested in the main patterns relatable to phenology rather than pixel-wise accuracy on the tiny fraction of the overall dataset where it would be feasible to produce extremely detailed validation data. We were also using GCC, a coarse image-level index for above-ground timeseries anyway and it was easier to produce a validation set without needing full or connective annotation of individual roots or complete accuracy in root diameters.

To process these validation images, we used Rhizovision Explorer (Seethepalli *et al*., 2021), a state-of-the-art root imagery tool. We processed the manual pixel maps to extract RSA and RLD without relying on the segmentation nor our post-processing scripts for RLD. Thus our validation of the RLD step is the most conservative option and differences could have arisen either from the trained CNN segmentation mismatch with manual annotation or the skeletonization mismatch with the software routine. In the analyses which followed, we used RLD for time series whenever possible as this ignored potential inconsistencies in root diameter which would disproportionately affect volume, i.e. RSA. This could have affected both segmentation and manual mark-up, and been artefactual or real. In comparison C efflux in E1 and for growth rates in E1 and E2 we used RSA. We made this latter choice because RSA is closer to C pools in biomass than RLD despite diameter stability uncertainty.

After considering images in E1 and E2, we realised that horizontal installation meant that images on the top of the tube were different than at the bottom. Thereafter, we treated the top of the tube as ‘truth’ and used this 3/8 of a complete rotational series for further analyses. In E3/4, where the instruments were deployed in a conventional fashion in the field, we used the sides of the instrument (excluding 1/4 of images at on the top and bottom of the tube) for analysis

### Data analysis

We took a daily average across all valid images for a single value which was not biased by any potential sub-daily cycle as an artefact of the segmentation (see Supplementary Fig. S1). For timeseries, we used a further three day rolling average in common with field above-ground phenology approaches (Migliavacca *et al*., 2011*a*; Aasen *et al*., 2020). To calculate growth rates in E1 and E2, we took the linear slope in the daily average over a five-day window. To analyse model accuracy and usefulness to replace manual annotation, we used reduced major axes regression in the *lmodel2* package (Legendre, 2018) in R (R Core Team, 2018). With this analysis we calculate determination coefficient and slope of observed (x, annotation) vs estimated (y, CNN RSA) in the imagery, accounting for similar magnitude of the error in x and y. Additionally, we quantified if systematic bias was introduced by confounding variables such as time, soil moisture, and absolute roots in the image by comparing these to the root length density from our method and checking for significant linear trends. We worked with normalized (0 to 1) timeseries to account for variation between mesocosms. To analyse the effect of mesocosm conditions on CO_2_ efflux in E1, we fit a GAM (generalized additive model (Hastie and Tibshirani, 1986; Wood, 2006), implemented via *mgcv* (Pedersen *et al*., 2019), which allowed an independent, non-linear smooth to be fit per predictor (e.g. a RSA effect on soil respiration could occur due to more biomass (positive slope) or turnover (negative slope)). For predictors, we used normalized (0 – 1) soil moisture content, temperature, 5-day slope of normalized RSA and normalized GCC and the mean normalized ‘biomass index’ (mean normalized GCC and normalized RSA, combined due to high concurvity). We fit a univariate smooth for each without interactions. Otherwise, concurvity in all cases was less than 0.7. We used the restricted maximum likelihood (REML) method to estimate smooths to reduce overfitting. We compared the variable importance in these models using the *varIMP* function in the *caret* package (Kuhn, 2008). For overall timeseries in E3 and E4, we gap-filled an aggregate timeseries level accounting for the occasional missing cycle via linear interpolation between successive points. The GAM analysis used only the non-gap filled datapoints.

## RESULTS

### Method Validation

We validated RSA and RLD from our simple extremely high throughput method against the state-of-the-art GUI based tools for the same property. Overall, we found a good match for RSA in E1 (R^2^ = 95%, Fig. 3A), E2 (R2 = 66%), E3 (R2 = 81 %), and E4 (R2 = 91 %). The relationship between manual and CNN root surface area for E2, E3 and E4 is shown in Supplementary Fig. S2. In the mesocosm experiments the CNN tended to identify more pixels as roots than humans (slope > 1), while in the field it was the reverse (slope < 1). We investigated the drivers of this disagreement in E1. While there was an expected relationship between manual cover and absolute difference between CNN and manual mark-up (bigger values could be more wrong, Fig. 3B), there was no effect of time, soil moisture content or manual cover on the error relative to the manual cover (Fig. 3C, 3D, 3E). Hence, we could trust the segmentation on average over the whole time series in terms of dynamics, and if adjusted by a linear transformation, in terms of magnitude.

**Fig. 3.**
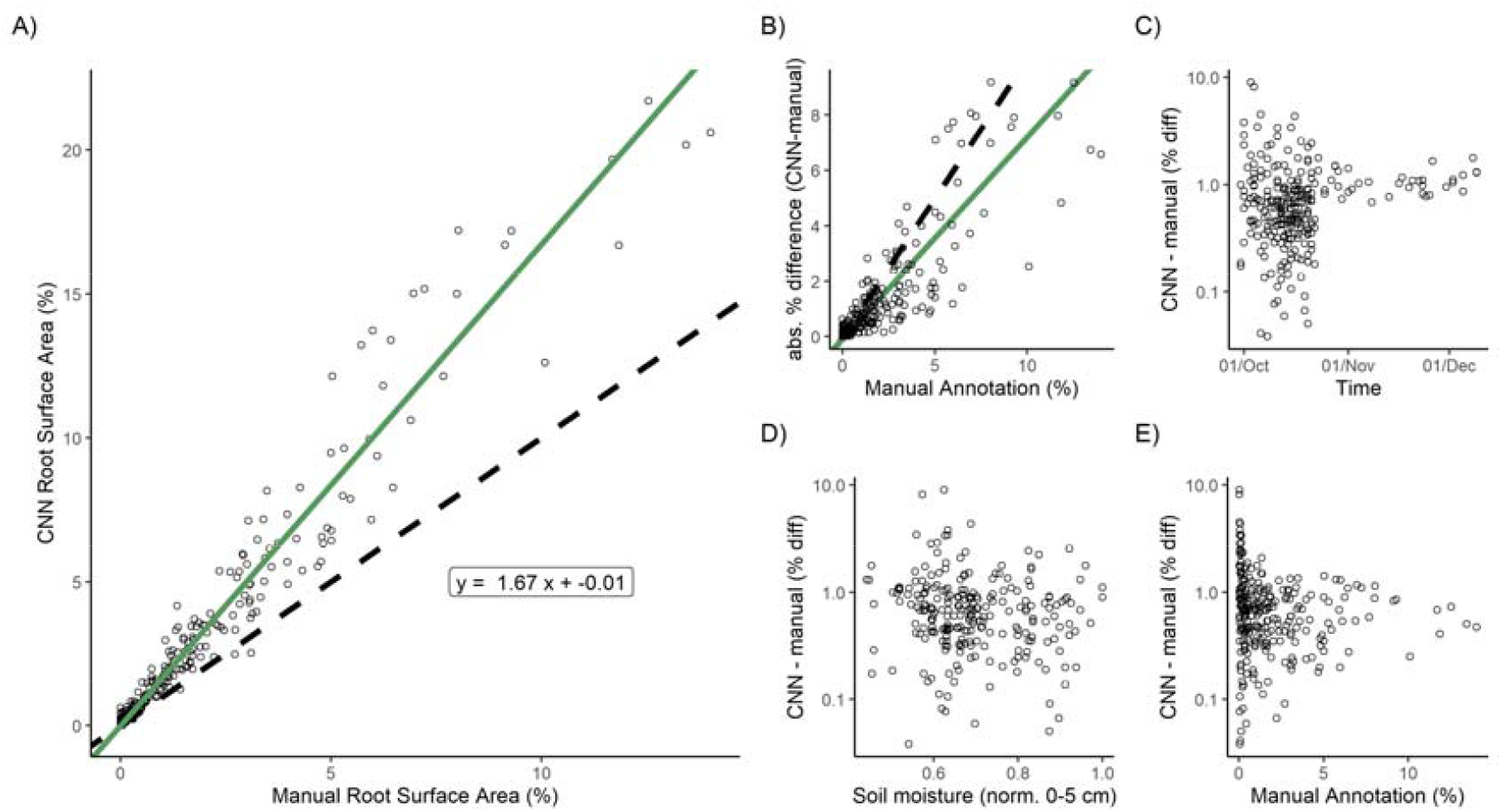
a) Validation of CNN-annotation for Root Surface Area (RSA) against independent manual root annotation in E1. The green line shows a reduced major axis regression of the two variables, while the dashed line is a 1:1 relationship. Statistics show equation of significant RMA fit. b) absolute percentage difference between manual RSA and CNN-classified RSA, and relative percent difference standardised to manual annotation over c) time, d) soil moisture content in the 0-5 cm soil at time of sampling and e) manual cover. The increased absolute error at higher manual cover was expected. The clustering of data towards the early part of panel c) is due to an uneven validation dataset.

We also found a good validation of RLD. While we used a simpler extraction routine with less tuneable parameters than the software, our RLD extraction agreed closely (Fig. 4). The R2 in E1 was 97%, in E2 was 68%, E3 was 84% and E4 87 %.

**Fig. 4.**
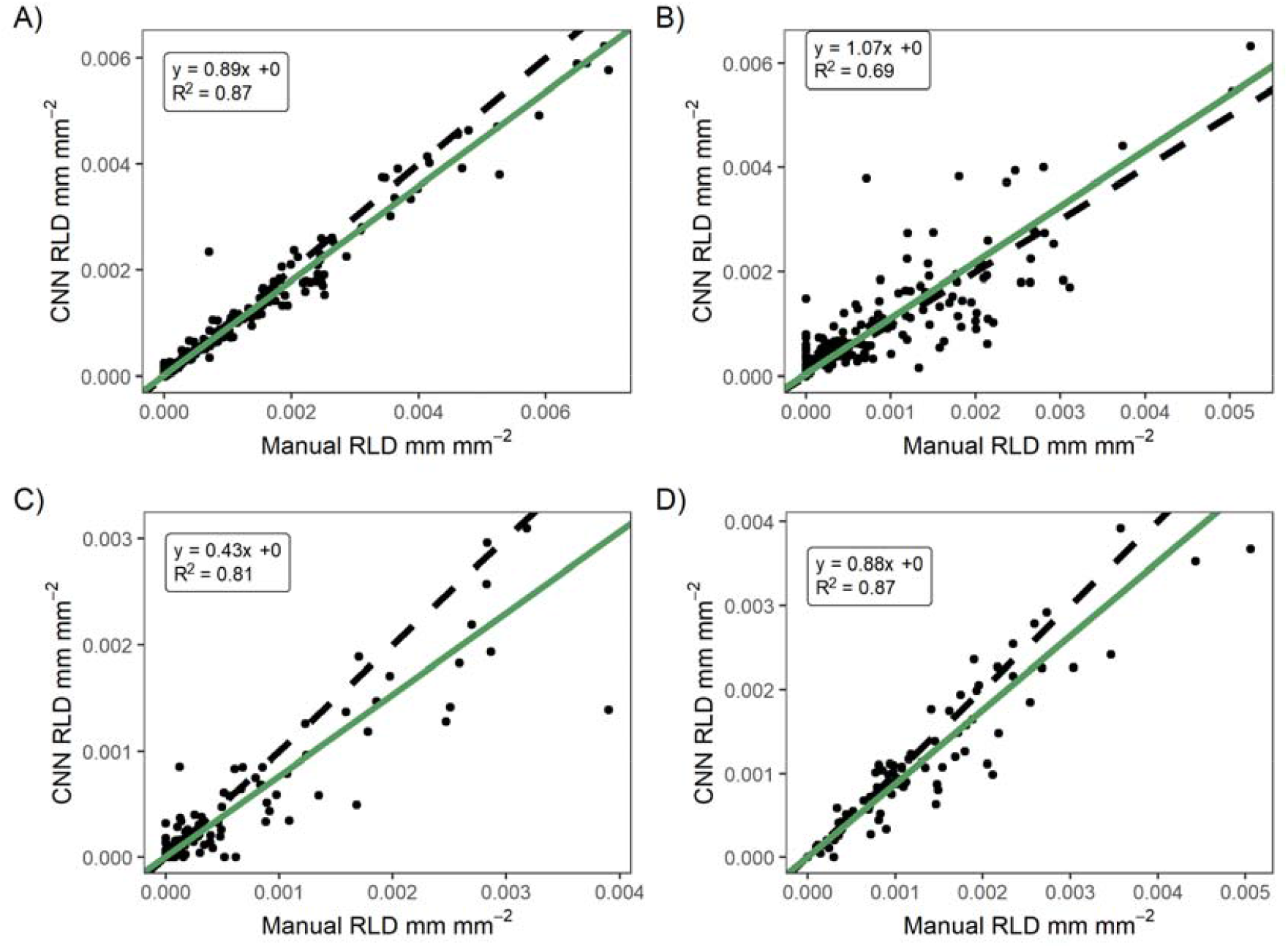
Validation of CNN-annotation for Root Length Density (RLD) using our simple script for the very large dataset against extraction of RLD on manual annotated imagery using the Rhizovision Explorer software tool. A is E1, B is E2, C is E3 and D is E4. In general there was good agreement, with disagreement due to both the accuracy of the CNN in segmenting (see Fig 2.) and potentially in differences in the RLD extraction routines.

### Experiment 1: Biological Interpretation

In E1 we paired the CNN-segmented RSA and RLD time series with the green chromatic coordinate (GCC) from PhenoCam imagery and the absolute root mass measurements (Fig. 5A). We found a good match between normalized root mass and RSA (Pearson r = 0.96), while RLD was smoother and slightly less well matched (Pearson r = 0.94), Some of the instability in root surface area appeared to follow the watering events (Fig 5A), a potential effect not caught by comparison with SWC nor visible in human mark-up.

**Fig. 5.**
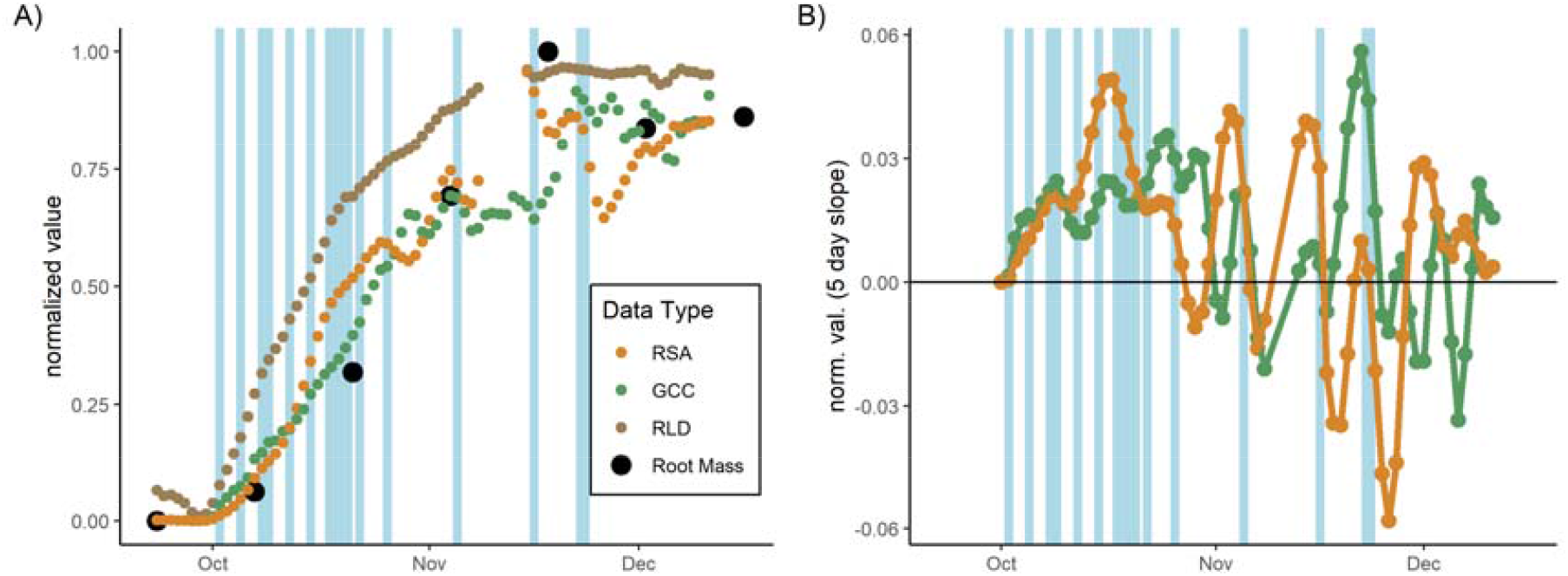
A time series of root data from the mesocosm experiment. Panel A) shows the relative, 3-day smoothed root surface area (RSA), root length density (RLD) GCC (greenness of above-ground vegetation), and root mass showing parallel development all scaled 0-1. Panel B) shows the 5-day slope of biomass-related values (i.e., rate of change) where at some periods GCC is increasing faster than root surface area and at others root surface area is increasing faster than GCC. Blue lines show watering events which are scaled in width relative to the volume of water

We also compared rates of change (‘growth’) of the above- and below-ground index conceptually closes to biomass. In general, this differed between GCC and RSA (Fig. 5B) except at the start of the experiment. As previously mentioned, some of this variation was due to the short period after watering, but this continued we ceased water inputs. In general, RSA growth continued to be positive at the end of the experiment even when the GCC was stable or decreasing as the soil slowly dried out.

Using the GAM on Experiment 1, we were able to explain 46% of the total variation in CO_2_ soil efflux with the best fitted model (Fig. 6A). In this model, soil moisture content (P < 0.001) and root growth (P < 0.001) were significant. Root growth was most important in determining soil CO_2_ efflux (Fig. 6B), increasing root growth rate had a positive effect on soil CO_2_ efflux (Fig. 6D). In contrast, absolute biomass, and GCC slopes, our proxy for leaf growth rate, did not have an effect (Fig 6C, 6E) on soil CO_2_ efflux.

**Fig. 6.**
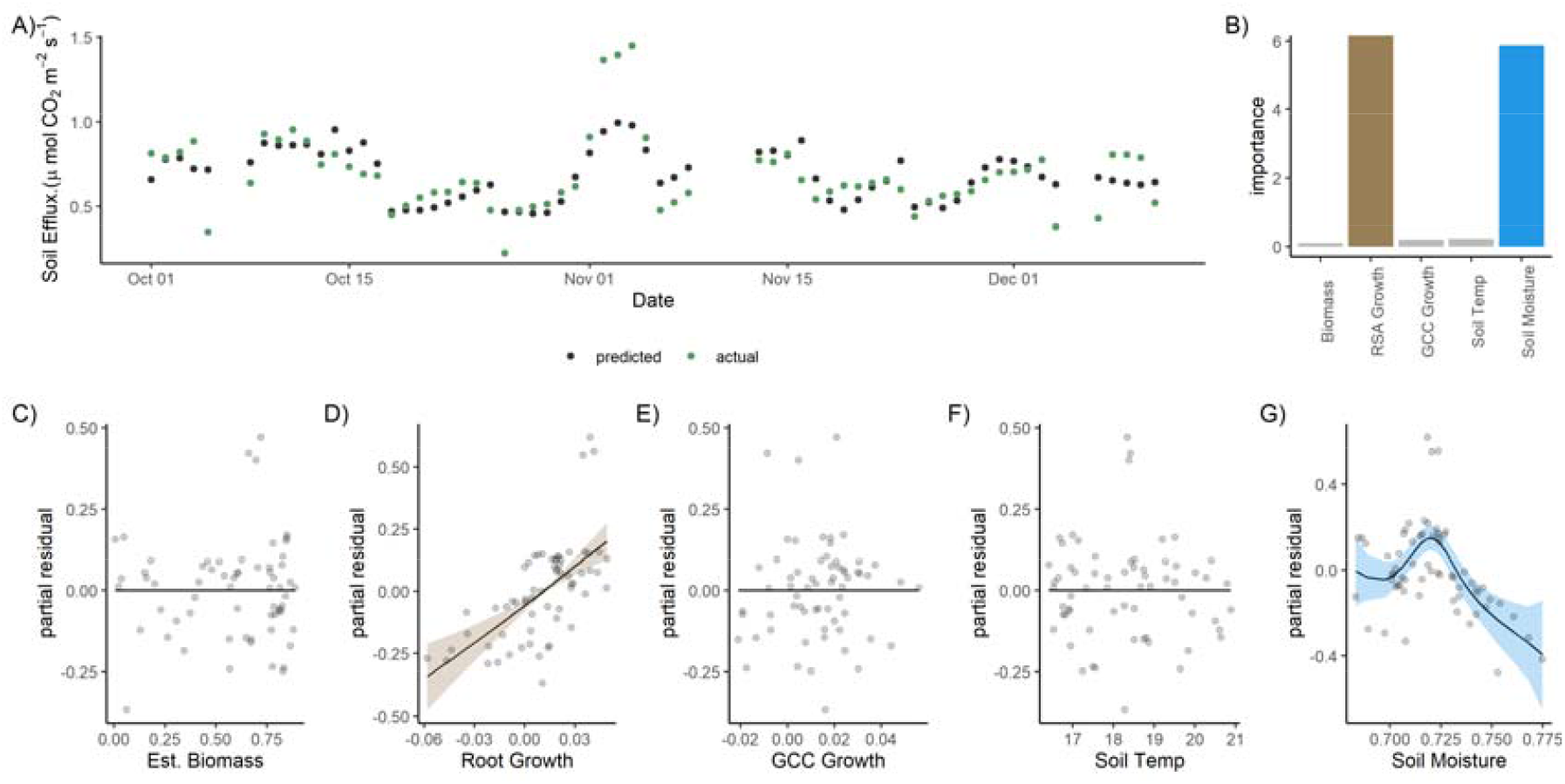
Summary of the GAM model fit for E1. Root surface area correlated with GCC so we used a mean of the two after normalization to represent biomass in the model (Est. Biomass). We were able to predict daily CO_2_ flux fairly well with an R^2^ of 46% (panel A). The model contained univariate smooths for Est. Biomass, shown vs partial residuals in panel C), the 5-day slopes (i.e. growth rate) of root surface area (panel D) and GCC (panel E), soil temperature (panel F) and soil moisture content (panel G). Shaded area shows 2 x SE. When variable importance was considered (panel b), the slope of root cover (i.e. change in amount of roots, birth or death) was a much better predictor than that of biomass.

### Experiment 2: Instrument Consistency

Overall, a similar time series was extractable from each mesocosm (Supplemental Fig. S3). Correlation between GCC and RLD was between 0.7 and 0.96. Notably, GCC increased faster than root surface area in the first four mesocosms (Fig. 7, Supplemental Fig. S3). The mesocosms were arranged in numerical order, suggesting that a gradient (e.g., light) within the greenhouse may have driven a difference. Once watering ceased, root growth rate declined less steeply than GCC in all mesocosms and remained positive for longer than GCC growth rate, indicating a continued production of roots even as the above-ground began to yellow.

**Fig. 7.**
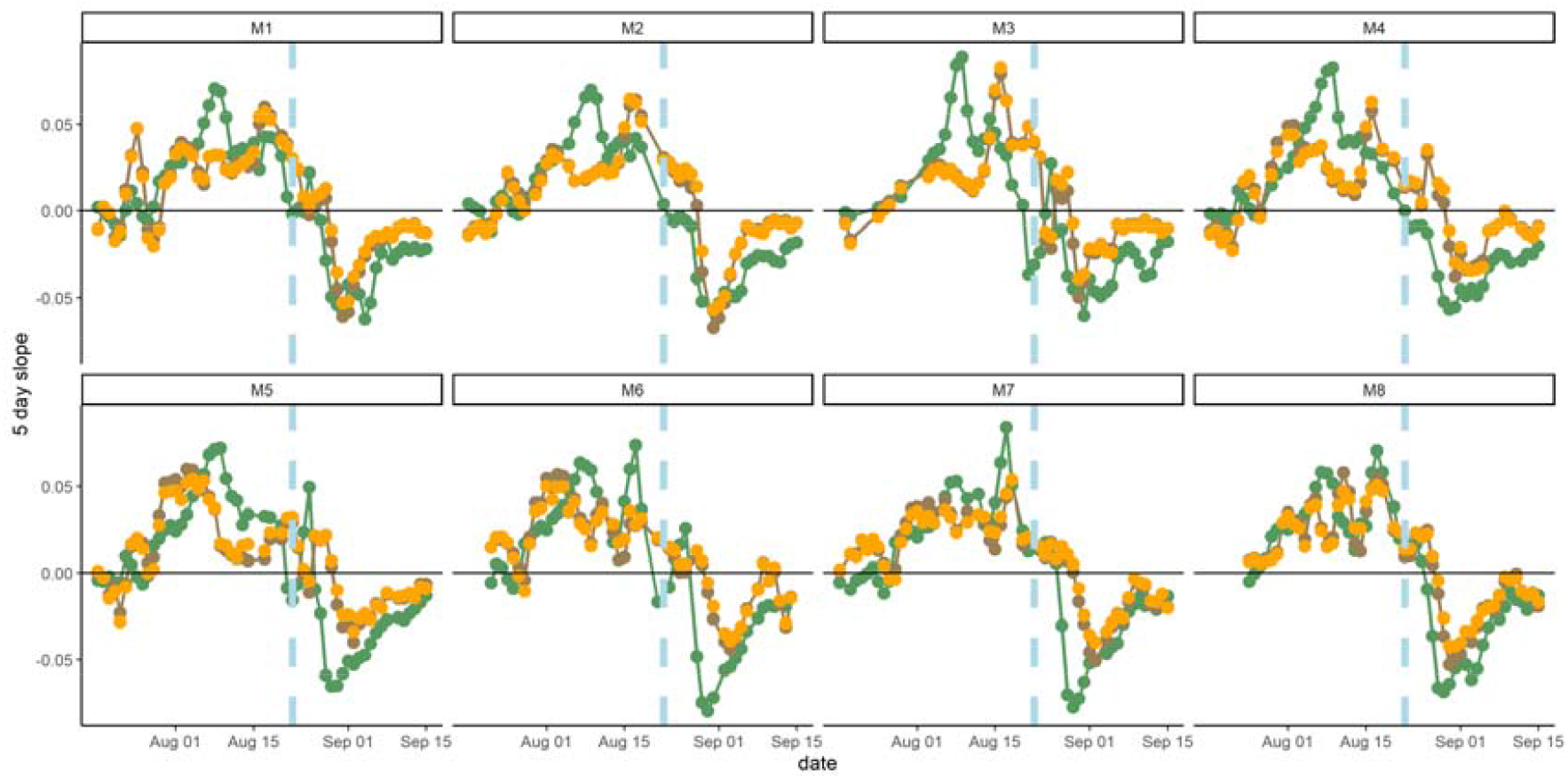
5-day rolling average slope (‘growth rate’), brown is RSA, orange is RLD, green is GCC across 8 mesocosms in E2. RLD and RSA are closely correlated. Vertical blue line indicates the last watering date, horizontal black line is 0 and the transition point between positive growth rates (above-) and a decrease in the index (below-) indicating yellowing or disappearance of roots identified by the CNN trained on ‘living’ roots.

The roots extracted from ingrowth cores at the start of the experiment matched the overall time series (when normalized between mesocosms, Supplemental Fig. S3B). However, the last data point, collected at harvest did not, and the measured root biomass was still high when RSA had dropped.

### Experiment 3: Root Litter Field Trial

The first field trial was conducted in a Mediterranean ecosystem so the period studied was early in the growing year. Examining mean total change in RLD over the whole depth and all instruments sampled, we observed growth period starting in November-December, and after the initial green-up of the above-ground system as detected by GCC (Fig. 8). Certain short-term instability in this index appeared to be related to periods of rainfall (as in E1). We additionally filtered out a short period (19-21 December) where all images across all instruments were very poorly illuminated. In general, temporal variability and noise in RLD was of similar relative magnitude to site-level GCC. RLD generally was lagged following positive GCC change, unlike E1, potentially due to the conventional installation angle here and subsequent better coverage of the whole soil. E3 was halted by the failure of accurate timekeeping across all instruments, which we subsequently addressed in E4.

**Fig. 8.**
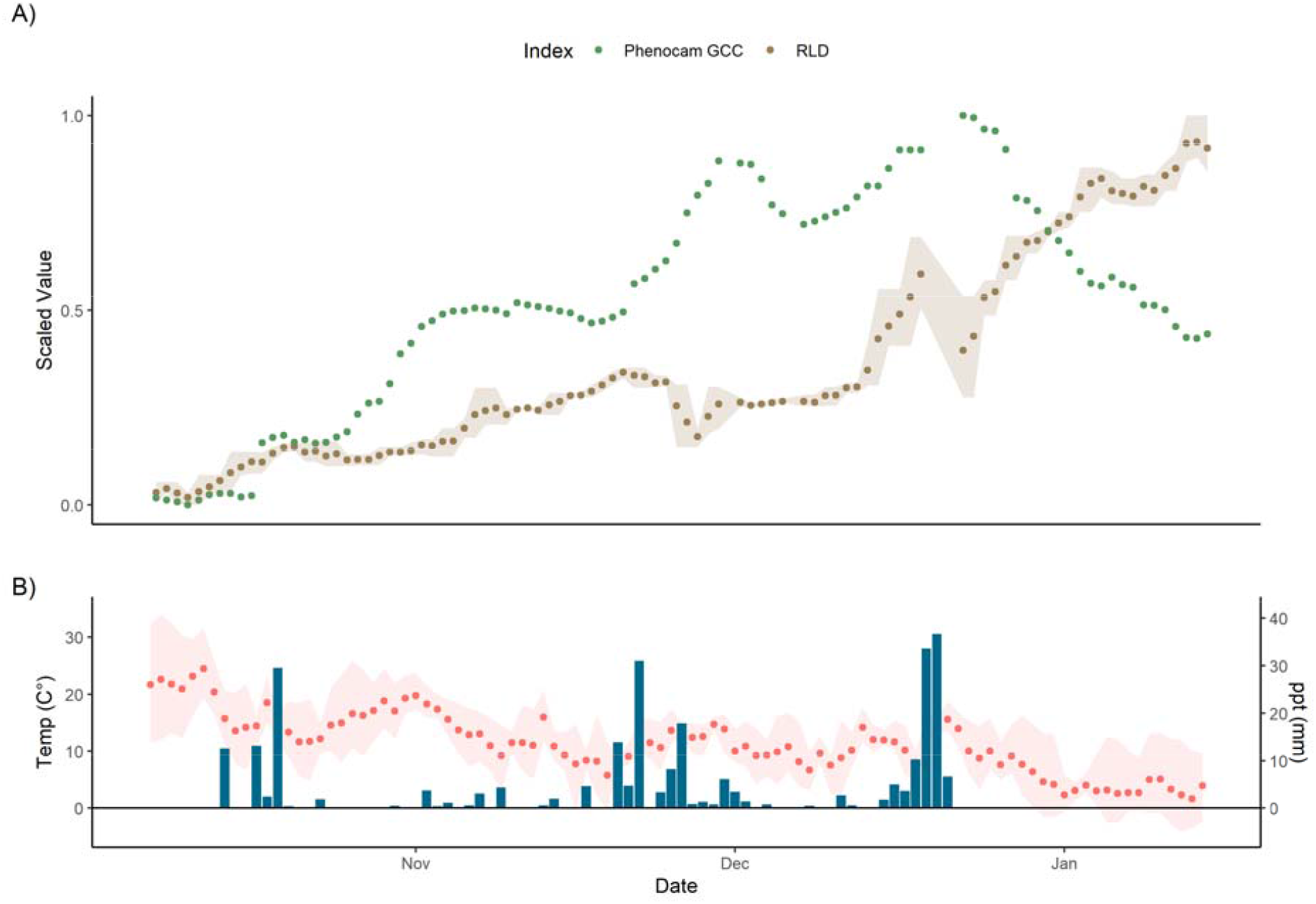
A) 3-day normalized GCC (green chromatic coordinate, green) and mean RLD (brown) from 3 instruments with at the Majadas de Tiétar site E Error bands on root cover correspond to maximum and minimum mean segmented root cover in the aggregation period. B) Air temperature and precipitation, taken as daily mean with error bands on temperature corresponding to observed maximum and minimum. The high errors in late November are due to instrument drop-out, reduction to two instruments in this period and subsequent instrument replacement, A short period (2 days) is removed in mid-December as batteries failed to charge in very cloudy weather. Root growth began later in the Mediterranean growing season than leaf growth and continued even when GCC was declining in midwinter. There was some instability in the root index which may have followed precipitation, although this was not larger than relative instability in GCC.

### Experiment 4: Low Temperature Field Trial

In E4 instruments ran without issue under winter conditions more severe than those which had previously caused timekeeping failures. We observed root growth through February and March (Fig. 9) with periodic increases in RLD and periods of no net growth which did not qualitatively appear to be linked to site conditions. We had occasional unexplained periods where the measurement cycle did not start; this affected ∼ 5 % of observational periods with no relationship to temperature or humidity. The images were confounded by condensation and soil animals, which we were largely able to successfully train around) so did not have the same vulnerability to soil moisture and precipitation as E1 and E3. There was no relationship between the root surface area we segmented over the whole 0-40 cm depth sampled and air temperature, precipitation, nor soil moisture. Hence the observations can be considered real root growth in this time.

**Fig. 9.**
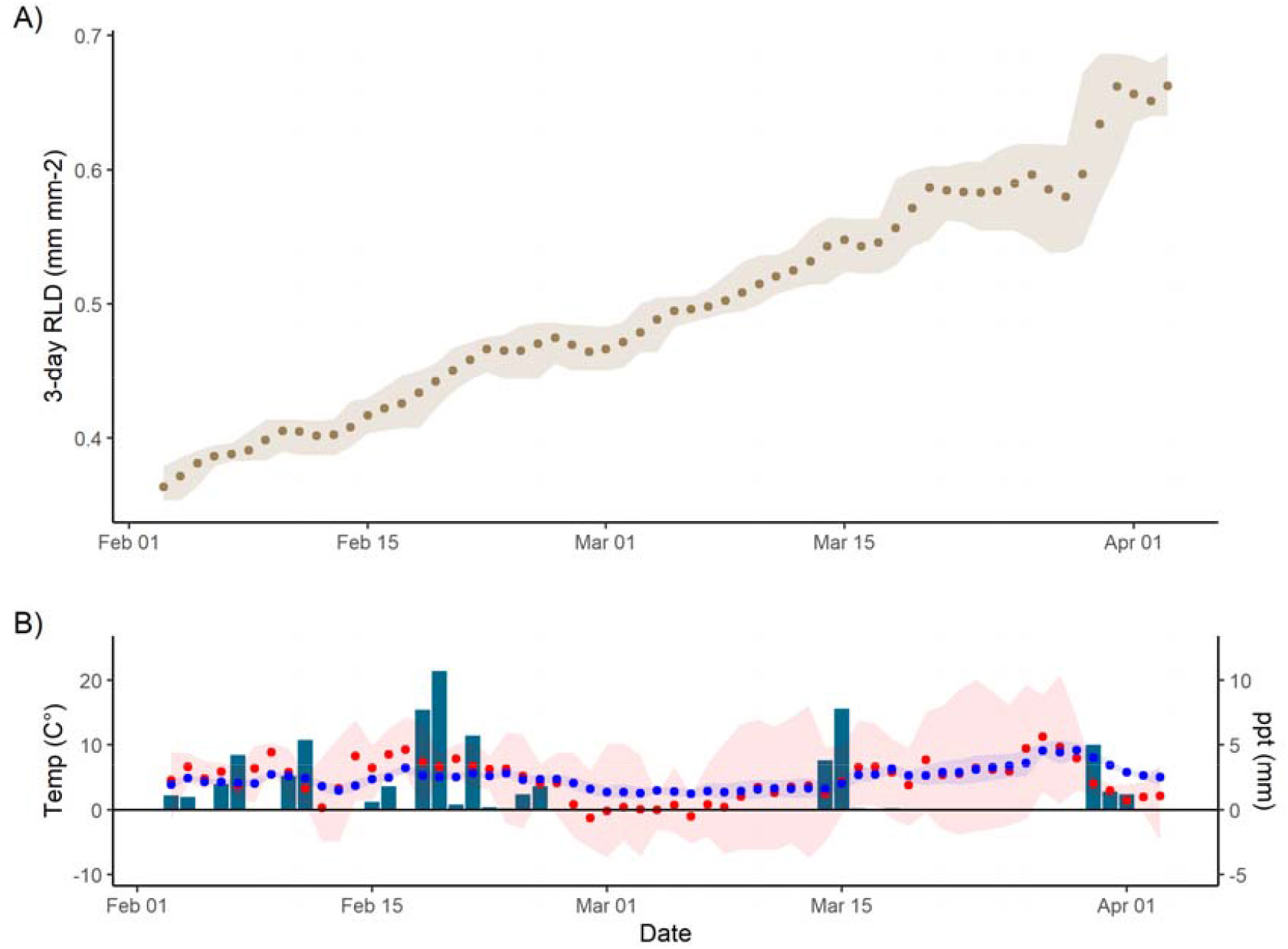
a) 3-Day mean normalized root length density (brown) from 2 instruments at the Jena Experiment site showing growth in February and March. Error band shows the maximum and minimum segmented RLD during the aggregation period. This winter trial shows robustness of the RMR to cold temperatures, the mean air temperature (red) and soil temperature at 8 cm (blue) in b) is shown with an error band corresponding to maximum and minimum daily temperatures which fell as low as -9°C. Unlike E3, it did not appear that in this experiment there was a sensitivity in root cover to precipitation.

## DISCUSSION

### Use of automatic minirhizotrons

The minirhizotron (MR) technique is the best method to measure root phenology and dynamics (Freschet *et al*., 2021). Automation of the whole workflow, from image collection to analysis, is essential for high time resolution data. Here, we presented a new automatic workflow. The system is designed for relatively affordable budgets (we used €2000 per instrument including development). We therefore replicate between instruments and can frequently resample through time. The method was robust in adverse field climatic conditions, after modification. We produced similar data from replicated instruments despite independent sensors (Fig. 7) for interpretable field time series (Fig. 8, Fig. 9). Methods such as PhenoCams (Sonnentag *et al*., 2012; Richardson *et al*., 2018) or networks for spectral vegetation indices (Gamon *et al*., 2015) are instrument standardised, but automatic minirhizotrons must combine robotics and imaging across multiple point sensors each with a short field of view. These technical aspects are also advancing fast. Soils are also complex and belowground imaging needs different magnification, sampling time, and power supplies depending on question. Hence, we recommend against standardization of every aspect of design if time and expertise is available to build to exact demand. In particular, cameras can be tooled to specific application, as neural networks likely require retraining to new sites and tasks and images must always be validated against properties of interest from manual annotation.

Replication is a key shortcoming of many field studies (Filazzola and Cahill Jr, 2021; Yang *et al*., 2022). These issues are especially acute for minirhizotrons given the short field of view, potential artefacts (Joslin and Wolfe, 1999) and huge range of soil and root appearances. The necessary modifications and COVID-19 led to field trials E3 and E4 not using all instruments from E2, but we observed consistency and reliability in patterns within instruments used. We also fully analysed the suitable data. While we discarded some of the interpreted images, we covered every sampling timepoint which is to our knowledge the numerically largest minirhizotron dataset so far analysed. Data collected at such high frequency have multiple issues even in this simple analysis with implications for application to more complex situations. The majority of this discussion concerns these issues.

### Interpreting High Resolution Root Image Data

Neural-network methods are an obvious approach to analysing minirhizotron images, like many other root phenotyping applications (Atkinson *et al*., 2019). They are also necessary to exploit the high volume of imagery from automated sampling. The main objective here was to pair existing methods with our automated sampling. While we did not develop a new CNN, we applied an established algorithm (Smith *et al*., 2020) for root images, because of the corrective annotation interface. This method performed well (Fig. 3, Fig. 4. Supplementary Fig. S2) despite the complexity of our images and the lack of time series consideration in design. We extracted RLD and RSA in an extremely simple fashion from segmented images, which contained noise but was compared with a similarly coarse above-ground index. More interpretive root phenotyping (Bauer *et al*., 2022) can also be paired to this CNN. In theory, many further architectural traits can be extracted with such a workflow. However, application to automatic measurements and field settings rely on high quality imagery, a CNN able to minimize artefacts in unpredictable circumstances and subjective parameter choice by the analyst, so we did not attempt this here. The large size of the automatic datasets also influenced our choice to process the segmented imagery ourselves and avoid using more sophisticated graphical user interface-based tools. The ability to run complex processing routines transparently from the command line would be a major advance for applying such methods to large automatic datasets and perform sensitivity analyses for tuning parameters in automated trait extraction.

Automatic root identification from automated sampling are demanding on training data but also on validation. We used annotated images produced for other CNN trials for validation data. These were human best practice and not objective truth so some difference between segmented and annotated images should be expected. Human annotations could also be outperformed in both consistency and accuracy alongside throughput in identifying true properties of interest by a sufficiently sophisticated neural network trained on good quality images. We also expected worse validation under field than greenhouse conditions because of the variety of soil and root appearances in the images. This was generally true, although the camera change for E2 and E3 meant that these validated less well than E1 and E4. (Fig. 3, Fig. 4, Supplementary Fig. S3). E2 was negatively affected by previous use of observatories (note the images in Fig. 4B with no manual RSA but a segmented area) and E3 followed a long fallow period so had many images with a ground truthing of no roots contributing to its *R*^*2*^. These ‘difficult’ images thus had multiple reasons for annotation to disagree with segmentation. In general, a suitable imaging sensor and attention to minimising observatory artefacts is critical for homemade minirhizotrons. However above-ground indexes are typically coarse in space and time (e.g. 3-day averaging PhenoCam data (Aasen *et al*., 2020)), so similar accuracy as E1 (95% RSA / 97% RLD) or E4 (91 % / 87 %) even E2 (66% / 68%) or E3 (81% / 84%%) with the less powerful camera is sufficient for generating plausible root time series (e.g. Fig 4, Fig S5, Fig 7,. Fig 8.). While in all four cases the CNN segmentation (plus simple processing for RLD) did not perfectly match 1:1 with manual RSA and RLD (Fig. 3, Fig 4, Supplementary Fig. S3) we were not validating nor training for true pixel-level identification. In any case, human root image interpretation is potentially biased by annotator and other methods such as soil core processing have their own artefacts. Considering some conversion between measurements always be necessary, consistent quantitative CNN interpretation has major advantages for throughput for root dynamics.

In the mesocosm E1, we also found a good agreement in physical validation between root mass and minirhizotron RSA (Fig. 5A, r = 0.96), and also in the growth period of Experiment 2 (but not the last, replicated measurement, discussed later). Minirhizotrons are not the best method for absolute biomass estimates at a single time, but further consideration is due to dynamic biomass estimates as the temporal resampling aspect cannot be achieved otherwise. There are differences in properties measurable from minirhizotrons and other methods (e.g. Addo-Danso et al., 2016; Milchunas, 2009; Nair et al., 2019), but minirhizotrons are a reasonable biomass index with appropriate conversions used (e.g. Brown et al., 2009; Johnson et al., 2001; Lee et al., 2017; Sullivan & Welker, 2005). Indeed, root identification relates directly to biological structures around a minirhizotron observatory. This can potentially be paired more easily with density and C contents than leaf greenness of a whole canopy, if the short field of view can be offset by replication. A key aspect of future studies will be to ascertain whether well-known artefacts of roots around minirhizotrons (Vamerali *et al*., 2009; Rytter and Rytter, 2012) can be understood to correct for these issues, and indeed if these artefacts vary with phenology or environmental conditions.

A further issue with our segmentation of high frequency images related to unrealistic variation after rain/watering (Fig. 5, Fig 8) and in E1 and E2, between different hours of the day (Supplementary Fig. S2). Like similar issues using classifiers for NIR-enabled minirhizotron images (Svane *et al*., 2019), sudden changes in soil colour/reflectance led to potential erroneous pixel identification and subsequent unrealistic changes in roots identified. We note that in Fig 8. this instability is not larger than similar short-term patterns in greenness above-ground, potentially due to illumination conditions. For the sub-daily artefact, this was not explainable by soil moisture perhaps because very local condensation at minirhizotron surfaces was not represented in the sensors. Indeed, sub-daily variation only became apparent after roots had colonized the sides of the observatories, suggesting that the CNN was misidentifying pixels in close proximity to roots. Inspection of the images confirms this explanation (Supplementary Fig. S1). Diel variation in root diameter (Huck *et al*., 1970) or hydraulic redistribution (passive movement of water via roots from wet to dry soil (Ryel, 2004)) at night and transpiration during the day drying these areas may provide an explanation which could bias a CNN more than human annotation. If this was a segmentation artefact which affected the immediate ‘rhizosphere’ only, this would not have been detected by our soil moisture sensors.

There are several potential solutions. The first is to aggregate, smooth or throw out data, which disregards information from high resolution sampling but is common practice in proximal remote sensing at ecosystem level (e.g. using smoothing splines, Migliavacca, Galvagno, et al., 2011). However, if one wants to analyse sub-daily data with potential diel patterns, this is not a viable solution unless using time series decomposition (e.g. Biriukova et al., 2021). Secondly, one could post-process with human intervention, applying an adjustment or a separately trained model, to periods with problematic changes. This interferes with the image index so is not favourable. Thirdly, one could train on more data, particularly around periods of difficulty. This exacerbates training data issues, especially if problematic events are rare but important. Fourthly, consistency in segmentation between sequential images could be used to filter for ‘true’ observations (i.e. basing interpretation on objective priors about root growth). Finally, a wholly different model structure incorporating other factors such as soil moisture or rainfall (i.e. describing when these issues could occur) or time series information (e.g. via recurrent network architectures such as Long-Short Term Memory approaches (Hochreiter and Schmidhuber, 1997), multivariate time series classification (Ruiz *et al*., 2021) or other time series classification (Fawaz *et al*., 2019)) could be built. This has the advantage of processing data without bias (if training data is selected fairly) at the cost of a more complex model and/or more variables to measure alongside root imagery.

### Field Robustness of Our Techniques

Both automation of measurement and image analysis are more advanced in simple artificial systems than field measurements, where litter, soil animals, hydrology and soil appearance complicate imagery. Instruments tested in controlled environments such as E1 and E2 and analysis methods also need to be robust in the field to study phenology. In the two short field studies we show here, we demonstrate viability of the technique. While E3 was compromised by camera quality and the structurally complex system, we produced a time series plausible from previous work at the site. In this ecosystem autumn root growth continues through winter, unlike above-ground vegetation indexes in most years (Nair *et al*., 2019). The instruments operated consistently on solar power for 4 months with 95 % uptime. While we do not show a whole phenological year nor include mass root death in the summer drought, this period contained undecomposed root litter following the arid summer, confusing for both human annotators and the CNN. Difficult minirhizotron images are troublesome for even experienced annotators (Peters *et al*., 2022) - a key advantage of an well trained automated approach is consistency. Our target was also a robust index comparable to above-ground digital repeat photography (e.g. Migliavacca, Galvagno, et al., 2011; Sonnentag et al., 2012) rather than exact match of segmentation to annotation. A key difference between minirhizotrons and proximal remote sensing is segmenting roots from soil and then interpreting this segmentation rather than a simple image index such as ‘greenness’ on a defined region of interest. Because we were segmenting complex features, minor variations in image setting may introduce short term variation. Future efforts could benchmark acceptable consistency for representativeness of phenology rather than requiring the same accuracy on a high-resolution dataset in a finer scale analysis from minirhizotrons. On the other hand, neural network approaches could be trained to *directly* interpret properties such as root length rather than segmenting then interpreting although this requires alternate models and data-demanding training if multiple traits are of interest.

E3 revealed an unexpected and random vulnerability of BIOS timekeeping to low temperatures which we corrected by adding a GPS clock. In E4 the system was able to run successfully at night-time temperatures as low as -7.5°C without issues. In this relatively ‘easy’ (stone-free and loamy) soil we were able to achieve a much close match with manual annotation. We could train around soil animals and condensation in this wet and cold part of the year and indeed found that roots were growing in this temperate winter. RSA almost doubled over the sixty days of the trial. Growth did not appear to be related to meteorological conditions, suggesting that i) these conditions were unlikely to be seriously biasing our CNN segmentation and ii) this early-season growth may be driven by intrinsic cues rather than immediate photosynthesis. Future longer timeseries will enable a deeper understanding of the coupled above-belowground action of the carbon cycle in plants. While we note our field indexes could be unstable, especially in E3, this was due to the low replication and differences between instruments rather than individual timeseries inconsistency. Wider application of such devices is hence reliant on reasonable per-instrument costs, the initial rationale for our instrument development.

### How Close Are We to True Automated Root Phenology Monitoring?

In comparison to all previous attempts to automate minirhizotron data collection and/or analysis, we produced high frequency timeseries in the lab and field (with support from an electrical/mechanical workshop and minimal computer science expertise). The workflow can process hundreds of images in the same time to annotate a single image conventionally and collecting images much more frequently than highest effort manual data collection. However, it does sacrifice some fidelity in extraction of root properties compared to manual methods. Many other properties (width, branching patterns *etc*.) may be extractable from an image compared to our two common architectural properties. These same trade-offs have already been made in leaf phenology monitoring (e.g. using ‘greenness’ or NDVI from remote sensing rather than counting leaves or measuring leaf angles). Minirhizotron imagery contains much more relevant structural information due to the extremely local scale and indeed these are regularly extracted manually. There is progress in doing this automatically (Seethepalli *et al*., 2021; Bauer *et al*., 2022) and we expect rapid advancement in future. Our analyses were also not particularly sensitive to roots overlapping or growing close due to overall low density. Moving from segmented images to full timeseries of multiple traits from multiple species in mixed communities and potentially dense root systems and tracking of individual root birth and death are ambitious and exciting goals which could be achieved by root ecologists and computer scientists collaborating in future. Indeed, similar tasks are possible in other biological image contexts (e.g. multilabel segmentation (Kubera et al., 2022), or dealing with overlapping via instance segmentation (Saleh et al., 2019)). Other CNN-minirhizotron applications can even tackle this last problem, albeit without the corrective annotation which made our approach useful for automatic data (Peters *et al*., 2022). Other possibilities such as root colour are readily available post-segmentation but we lack a theoretical understanding of the meaning of such timeseries in field communities where individual species differ alongside change in time. In terms of general trends (but not absolute mass), our RSA was also a reasonable proxy for root biomass so with appropriate calibration and density/area parameterisation imagery could be interpreted as changes in C pools and provide data to inform allocation in vegetation models. Interestingly the ‘harvest’ data point in E2 did not match the minirhizotron index, even when scaled. This is potentially explainable by the CNN, trained on live roots, outperforming humans sorting physical samples who also make mistakes in living/dead identification. Our datasets contained a period of time where there was litter in the field (E3) but not a specific period of turnover which may be more difficult for CNNs although roots did disappear from individual timeseries in all experiments. Because the living/dead distinction is of functional interest, further development e.g. training models specifically on dead roots, using multi-classifier segmentation or generating labels using the near-infrared or other multispectral images (Arnold *et al*., 2017; Bodner *et al*., 2017) and segmenting using RGB images may allow a segmentation of complex field imagery necessary for precise quantification of living root biomass and its complex and dynamic contribution to ecosystem C cycling.

A major advantage of the minirhizotron method is resampling over long timescales in the field. With this information, root:leaf asynchrony (Steinaker and Wilson, 2008; Steinaker *et al*., 2010; Sloan *et al*., 2016) can be understood from data (e.g. E1, E2, E3). Links to carbon cycling (e.g. E1) such as between photosynthesis, growth, and soil/ecosystem respiration (Bahn *et al*., 2008, 2009; Migliavacca *et al*., 2011*b*) could also be understood in scalable contexts. We were not able to make field observations of this duration in this study and long-term reliability remains the biggest unknown. Nonetheless we did target the most ‘difficult’ times of the year in E3 and E4 where cold and wet conditions could affect instrument performance. The accelerated clocks used in E1 and E2 also partially demonstrate long term performance which could capture whole annual cycles if performed at similar rates in the field as in E3. Minirhizotron studies may also use huge replication – in agricultural settings this can reach thousands of observatories (Rajurkar *et al*., 2022). In this case no matter how affordable at budget an automated system is, entire coverage is unlikely to be possible with robotic systems. For this we recommend pairing automated systems (to capture temporal dynamics) with manual systems (for spatial dynamics), ideally with exactly the same camera setup and processing workflow. In such a design automated phenology data can complement the most informative sampling times and spatial understanding can inform the most representative sites for scalable temporal trends.

## ABBREVIATIONS

CNN: Convolutional Neural Network
GCC: Green Chromatic Coordinate
NDVI: Normalized Difference Vegetation Index
RLD: Root Length Density
ROI: Region of Interest
RSA: Root Surface Area
RMR: Robotic Minirhizotron (the instrument developed in this study)

## SUPPLEMENTARY DATA

Supplementary data to this manuscript is available.

Supplementary Protocol S1: Full Technical Instrument Design

Supplementary Fig. S1: Example of Sub-Daily Diameter Artefacts found in mesocosm

Supplementary Fig. S2: Validation of RSA and RLD in E2, E3, E4.

Supplementary Fig. S3: Comparison between mesocosms in E2.

## ACKNOWLEDGEMENTS

This project has received funding from the European Union’s Horizon 2020 research and innovation programme under the Marie Sklodowska-Curie grant agreement No. 748893 (Marie Sklodowska-Curie Individual Fellowship to R. Nair). This research also received funding through the Alexander von Humboldt Foundation and the Max Planck Research Prize 2013 to M. Reichstein. VR was supported by the regional government of Extremadura (Spain) through a “Talento” fellowship (TA18022). We also thank Y. Luo for processing the PhenoCam-GCC data at MDT, A. Smith for development and user-friendly implementation of the *Rootpainter* CNN, M.Schrumpf for her very useful comments on this manuscript and M. Reichstein for his support for the project in general.

## AUTHORS CONTRIBUTIONS

Conceptualization: RN, MM. Methodology: RN, MS, MH, OK.: RN. Investigation: RN, VR. Data Curation, Formal Analysis, Software: RN. Writing – original draft: RN, Writing – review & editing: MM, VR

## FUNDING

### DATA AVAILABILITY

A dataset with validation data used in this study, along with a small dataset of unannotated images will be uploaded to Zenodo.org on final publication. The full image dataset is not sharable using current services due to its size, but the authors are happy to share this to developers of root segmentation tools on request. The code which we used to extract root length from segmented images is also available.

## REFERENCES

Aasen H, Kirchgessner N, Walter A, Liebisch F. 2020. PhenoCams for Field Phenotyping: Using Very High Temporal Resolution Digital Repeated Photography to Investigate Interactions of Growth, Phenology, and Harvest Traits. Frontiers in Plant Science 11, 593.

Abramoff RZ, Finzi AC. 2015. Are above- and below-ground phenology in sync? New Phytologist 205, 1054–1061.

Adair KL, Lindgreen S, Poole AM, Young LM, Bernard-Verdier M, Wardle DA, Tylianakis JM. 2019. Above and belowground community strategies respond to different global change drivers. Scientific Reports 9, 2540.

Addo-Danso SD, Prescott CE, Smith AR. 2016. Methods for Estimating Root Biomass and Production in Forest and Woodland Ecosystem Carbon Studies: A Review. Forest Ecology and Management 359, 332–351.

Allen MF, Kitajima K. 2013. In Situ High-Frequency Observations of Mycorrhizas. New Phytologist 200, 222–228.

Allen MF, Kitajima K. 2014. Net Primary Production of Ectomycorrhizas in a California Forest. Fungal Ecology 10, 81–90.

Allen MF, Vargas R, Eric A. Graham, et al. 2007. Soil Sensor Technology: Life within a Pixel. BioScience 57, 859.

Alonso-Crespo IM, Weidlich EWA, Temperton VM, Delory BM. 2022. Assembly history modulates vertical root distribution in a grassland experiment. Oikos n/a, e08886.

Arnold T, Leitner R, Bodner G. 2017. Near infrared hyperspectral imaging system for root phenotyping. Sensing for Agriculture and Food Quality and Safety IX. SPIE, 94–99.

Atkinson JA, Pound MP, Bennett MJ, Wells DM. 2019. Uncovering the hidden half of plants using new advances in root phenotyping. Current Opinion in Biotechnology 55, 1–8.

Bahn M, Rodeghiero M, Anderson-Dunn M, et al. 2008. Soil Respiration in European Grasslands in Relation to Climate and Assimilate Supply. Ecosystems 11, 1352–1367.

Bahn M, Schmitt M, Siegwolf R, Richter A, Brüggemann N. 2009. Does photosynthesis affect grassland soil-respired CO2 and its carbon isotope composition on a diurnal timescale? New Phytologist 182, 451–460.

Bauer FM, Lärm L, Morandage S, Lobet G, Vanderborght J, Vereecken H, Schnepf A. 2022. Development and Validation of a Deep Learning Based Automated Minirhizotron Image Analysis Pipeline. Plant Phenomics 2022.

Beluch WH, Genewein T, Nürnberger A, Köhler JM. 2018. The Power of Ensembles for Active Learning in Image Classification. 9368–9377.

Biriukova K, Pacheco-Labrador J, Migliavacca M, Mahecha MD, Gonzalez-Cascon R, Martín MP, Rossini M. 2021. Performance of Singular Spectrum Analysis in Separating Seasonal and Fast Physiological Dynamics of Solar-Induced Chlorophyll Fluorescence and PRI Optical Signals. Journal of Geophysical Research: Biogeosciences 126, e2020JG006158.

Blume-Werry G, Jansson R, Milbau A. 2017. Root phenology unresponsive to earlier snowmelt despite advanced above-ground phenology in two subarctic plant communities. Functional Ecology 31, 1493–1502.

Bodner G, Alsalem M, Nakhforoosh A, Arnold T, Leitner D. 2017. RGB and Spectral Root Imaging for Plant Phenotyping and Physiological Research: Experimental Setup and Imaging Protocols. Journal of Visualized Experimentslll: JoVE, 56251.

Brown ALP, Day FP, Stover DB. 2009. Fine Root Biomass Estimates from Minirhizotron Imagery in a Shrub Ecosystem Exposed to Elevated CO2. Plant and Soil 317, 145–153.

Budd S, Robinson EC, Kainz B. 2021. A survey on active learning and human-in-the-loop deep learning for medical image analysis. Medical Image Analysis 71, 102062.

De Kauwe MG, Medlyn BE, Zaehle S, et al. 2014. Where Does the Carbon Go? A Model-Data Intercomparison of Vegetation Carbon Allocation and Turnover Processes at Two Temperate Forest Free-Air CO2 Enrichment Sites. New Phytologist 203, 883–99.

Defrenne CE, Childs J, Fernandez CW, Taggart M, Nettles WR, Allen MF, Hanson PJ, Iversen CM. 2021. High-resolution minirhizotrons advance our understanding of root-fungal dynamics in an experimentally warmed peatland. Plants, People, Planet 3, 640–652.

Delory BM, Baudson C, Brostaux Y, Lobet G, du Jardin P, Pagès L, Delaplace P. 2016. archiDART: An R Package for the Automated Computation of Plant Root Architectural Traits. Plant and Soil 398, 351–365.

Dijkstra FA, Zhu B, Cheng W. 2021. Root effects on soil organic carbon: a double-edged sword. New Phytologist 230, 60–65.

El-Madany TS, Reichstein M, Perez-Priego O, et al. 2018. Drivers of Spatio-Temporal Variability of Carbon Dioxide and Energy Fluxes in a Mediterranean Savanna Ecosystem. Agricultural and Forest Meteorology 262, 258–278.

Fawaz HI, Forestier G, Weber J, Idoumghar L, Muller P-A. 2019. Deep learning for time series classification: a review. Data Mining and Knowledge Discovery 33, 917–963.

Filazzola A, Cahill Jr JF. 2021. Replication in field ecology: Identifying challenges and proposing solutions. Methods in Ecology and Evolution 12, 1780–1792.

Freschet GT, Pagès L, Iversen CM, et al. 2021. A starting guide to root ecology: strengthening ecological concepts and standardising root classification, sampling, processing and trait measurements. New Phytologist 232, 273–1122

Gamon JA, Kovalchuck O, Wong CYS, Harris A, Garrity SR. 2015. Monitoring seasonal and diurnal changes in photosynthetic pigments with automated PRI and NDVI sensors. Biogeosciences 12, 4149–4159.

Gillert A, Peters B, Freiherr von Lukas, U, Kreyling, J. 2021. Identification and Measurement of Individual Roots in Minirhizotron Images of Dense Root Systems. Proceedings of the IEEE/CVF International Conference on Computer Vision (ICCV) Workshops. 1323–1331.

Han E, Smith AG, Kemper R, White R, Kirkegaard JA, Thorup-Kristensen K, Athmann M. 2021. Digging roots is easier with AI. Journal of Experimental Botany 72, 4680–4690.

Hastie T, Tibshirani R. 1986. Generalized Additive Models. Statistical Science 1, 297–310.

Herrmann S, Grams TEE, Tarkka MT, et al. 2016. Endogenous rhythmic growth, a trait suitable for the study of interplays between multitrophic interactions and tree development. Perspectives in Plant Ecology, Evolution and Systematics 19, 40–48.

Hochreiter S, Schmidhuber J. 1997. Long Short-Term Memory. Neural Computation 9, 1735–1780.

Huck MG, Klepper B, Taylor HM. 1970. Diurnal Variations in Root Diameter. Plant Physiology 45, 529–530.

Huo C, Cheng W. 2019. Improved root turnover assessment using field scanning rhizotrons with branch order analysis. Ecosphere 10, e02793.

Iversen CM, Murphy MT, Allen MF, Childs J, Eissenstat DM, a. Lilleskov E, Sarjala TM, Sloan VL, Sullivan PF. 2011. Advancing the Use of Minirhizotrons in Wetlands. Plant and Soil 352, 23–39.

Johnson MG, Tingey DT, Phillips DL, Storm MJ. 2001. Advancing fine root research with minirhizotrons. Environmental and Experimental Botany 45, 263–289.

Joslin JD, Wolfe MH. 1999. Disturbances During Minirhizotron Installation Can Affect Root Observation Data. Soil Science Society of America Journal 63, 218–221.

Kubera E, Kubik-Komar A, Kurasiński P, Piotrowska-Weryszko K, Skrzypiec M. 2022. Detection and Recognition of Pollen Grains in Multilabel Microscopic Images. Sensors 22, 2690.

Kuhn M. 2008. Building Predictive Models in R Using the caret Package. Journal of Statistical Software 28, 1–26.

Le Marié C, Kirchgessner N, Marschall D, Walter A, Hund A. 2014. Rhizoslides: Paper-Based Growth System for Non-Destructive, High Throughput Phenotyping of Root Development by Means of Image Analysis. Plant methods 10, 13.

Lee CG, Suzuki S, Noguchi K, Inubushi K. 2017. Estimation of Fine Root Biomass Using a Minirhizotron Technique among Three Vegetation Types in a Cool-Temperate Brackish Marsh. Soil Science and Plant Nutrition 62, 465–470.

Legendre P. 2018. R package ‘lmodel2’ v 1.7-3. https://cran.r-project.org/web/packages/lmodel2/index.html Accessed 01/06/2022

Liu S, Barrow CS, Hanlon M, Lynch JP, Bucksch A. 2021. DIRT/3D: 3D root phenotyping for field-grown maize (Zea mays). Plant Physiology 187, 739–757.

Luo Y, El-Madany TS, Filippa G, Ma X, Ahrens B, Carrara A, Gonzalez-cascon R, Cremonese E, Galvagno M, Tiana W. 2018. Using Near-Infrared-Enabled Digital Repeat Photography to Track Structural and Physiological Phenology in Mediterranean Tree – Grass Ecosystems. Remote Sensing 10, 1293.

Luo Y, El-Madany T, Ma X, et al. 2020. Nutrients and water availability constrain the seasonality of vegetation activity in a Mediterranean ecosystem. Global Change Biology 26, 4379–4400.

Migliavacca M, Galvagno M, Cremonese E, et al. 2011a. Using digital repeat photography and eddy covariance data to model grassland phenology and photosynthetic CO2 uptake. Agricultural and Forest Meteorology 151, 1325–1337.

Migliavacca M, Reichstein M, Richardson AD, et al. 2011b. Semiempirical modeling of abiotic and biotic factors controlling ecosystem respiration across eddy covariance sites. Global Change Biology 17, 390–409.

Milchunas DG. 2009. Estimating Root Production: Comparison of 11 Methods in Shortgrass Steppe and Review of Biases. Ecosystems 12, 1381–1402.

Nagel KA, Putz A, Gilmer F, et al. 2012. GROWSCREEN-Rhizo Is a Novel Phenotyping Robot Enabling Simultaneous Measurements of Root and Shoot Growth for Plants Grown in Soil-Filled Rhizotrons. Functional Plant Biology 39, 891–904.

Nair RKF, Morris KA, Hertel M, Luo Y, Moreno G, Reichstein M, Schrumpf M, Migliavacca M. 2019. N:P stoichiometry and habitat effects on Mediterranean savanna seasonal root dynamics. Biogeosciences 16, 1883–1901.

Nijland W, de Jong R, de Jong SM, Wulder MA, Bater CW, Coops NC. 2014. Monitoring plant condition and phenology using infrared sensitive consumer grade digital cameras. Agricultural and Forest Meteorology 184, 98–106.

Pedersen EJ, Miller DL, Simpson GL, Ross N. 2019. Hierarchical generalized additive models in ecology: an introduction with mgcv. PeerJ 7, e6876.

Peters B, Blume-Werry G, Gillert A, Schwieger S, von Lukas UF, Kreyling J. 2022. As good as human experts in detecting plant roots in minirhizotron images but efficient and reproducible: The Convolutional Neural Network “RootDetector”. https://doi.org/10.21203/rs.3.rs-1434431/v1 [preprint]

Poorter H, Bühler J, van Dusschoten D, Jos Climent Postma, Johannes A. 2012. Pot size matters: A meta-analysis of the effects of rooting volume on plant growth. Functional Plant Biology 39, 839–850.

R Core Team. 2018. R: A Language and Environment for Statistical Computing. Vienna, Austria. R Foundation for Statistical Computing https://www.R-project.org/.

Radville L, McCormack ML, Post E, Eissenstat DM. 2016. Root Phenology in a Changing Climate. Journal of Experimental Botany 67, erw062..

Rahmanzadeh H, Shojaedini SV. 2016. Novel Automated Method for Minirhizotron Image Analysislll: Root Detection Using Curvelet Transform. 29, 337–346.

Rajurkar AB, McCoy SM, Ruhter J, Mulcrone J, Freyfogle L, Leakey ADB. 2022. Installation and imaging of thousands of minirhizotrons to phenotype root systems of field-grown plants. Plant Methods 18, 39.

Raupach MR, Canadell JG, Ciais P, Friedlingstein P, Rayner PJ, Trudinger CM. 2011. The Relationship between Peak Warming and Cumulative CO2 Emissions, and Its Use to Quantify Vulnerabilities in the Carbon-Climate-Human System. Tellus, Series B: Chemical and Physical Meteorology 63, 145–164.

Ren P, Xiao Y, Chang X, Huang P-Y, Li Z, Gupta BB, Chen X, Wang X. 2021. A Survey of Deep Active Learning. ACM Computing Surveys 54, 180:1-180:40.

Richardson AD, Hufkens K, Milliman T, et al. 2018. Tracking Vegetation Phenology across Diverse North American Biomes Using PhenoCam Imagery. Scientific Data 5, 1–24.

Richardson AD, Keenan TF, Migliavacca M, Ryu Y, Sonnentag O, Toomey M. 2013. Climate change, phenology, and phenological control of vegetation feedbacks to the climate system. Agricultural and Forest Meteorology 169, 156–173.

Roscher C, Schumacher J, Baade J, Wilcke W, Gleixner G, Weisser WW, Schmid B, Schulze E-D. 2004. The role of biodiversity for element cycling and trophic interactions: an experimental approach in a grassland community. Basic and Applied Ecology 5, 107–121.

Ruiz AP, Flynn M, Large J, Middlehurst M, Bagnall A. 2021. The great multivariate time series classification bake off: a review and experimental evaluation of recent algorithmic advances. Data Mining and Knowledge Discovery 35, 401–449.

Ryel RJ. 2004. Hydraulic Redistribution. In: Esser K,, In: Lüttge u,, In: Beyschlag W,, In: Murata J, eds. Progress in Botany. Progress in Botany: Genetics Physiology Systematics Ecology. Berlin, Heidelberg: Springer, 413–435.

Rytter R-M, Rytter L. 2012. Quantitative estimates of root densities at minirhizotrons differ from those in the bulk soil. Plant and Soil 350, 205–220.

Saleh HM, Saad NH, Isa NAM. 2019. Overlapping Chromosome Segmentation using U-Net: Convolutional Networks with Test Time Augmentation. Procedia Computer Science 159, 524–533.

Seethepalli A, Dhakal K, Griffiths M, Guo H, Freschet GT, York LM. 2021. RhizoVision Explorer: open-source software for root image analysis and measurement standardization. AoB PLANTS 13.

Sloan VL, Fletcher BJ, Phoenix GK. 2016. Contrasting synchrony in root and leaf phenology across multiple sub-Arctic plant communities. Journal of Ecology 104, 239–248.

Smith AG, Han E, Petersen J, Olsen NAF, Giese C, Athmann M, Dresbøll DB, Thorup-Kristensen K. 2020. RootPainter: Deep Learning Segmentation of Biological Images with Corrective Annotation. bioRxiv, https://doi.org/10.1101/2020.04.16.044461 [preprint]

Sonnentag O, Hufkens K, Teshera-Sterne C, Young AM, Friedl M, Braswell BH, Milliman T, O’Keefe J, Richardson AD. 2012. Digital Repeat Photography for Phenological Research in Forest Ecosystems. Agricultural and Forest Meteorology 152, 159–177.

Steinaker DF, Wilson SD. 2008. Phenology of Fine Roots and Leaves in Forest and Grassland. Journal of Ecology 96, 1222–1229.

Steinaker DF, Wilson SD, Peltzer DA. 2010. Asynchronicity in Root and Shoot Phenology in Grasses and Woody Plants. Global Change Biology 16, 2241–2251.

Sullivan PF, Welker JM. 2005. Warming chambers stimulate early season growth of an arctic sedge: results of a minirhizotron field study. Oecologia 142, 616–626.

Svane SF, Dam EB, Carstensen JM, Thorup-Kristensen K. 2019. A multispectral camera system for automated minirhizotron image analysis. Plant and Soil 441, 657–672.

The GIMP Development Team. 2019. GIMP.

Vamerali T, Ganis A, Mosca G. 2009. Methods For Thresholding Minirhizotron Root Images., 1–2.

Vargas R, Allen MF. 2008. Dynamics of Fine Root, Fungal Rhizomorphs, and Soil Respiration in a Mixed Temperate Forest: Integrating Sensors and Observations. Vadose Zone Journal 7, 1055.

Vincent C, Rowland D, Na C, Schaffer B. 2016. A High-Throughput Method to Quantify Root Hair Area in Digital Images Taken in Situ. Plant and Soil.

Walker AP, Zaehle S, Medlyn BE, et al. 2015. Predicting long-term carbon sequestration in response to CO2 enrichment: How and why do current ecosystem models differ? Global Biogeochemical Cycles 29, 476–495.

Wang T, Rostamza M, Song Z, Wang L, McNickle G, Iyer-Pascuzzi AS, Qiu Z, Jin J. 2019. SegRoot: A high throughput segmentation method for root image analysis. Computers and Electronics in Agriculture 162, 845–854.

Weisser WW, Roscher C, Meyer ST, et al. 2017. Biodiversity effects on ecosystem functioning in a 15-year grassland experiment: Patterns, mechanisms, and open questions. Basic and Applied Ecology 23, 1–73.

Wood SN. 2006. Generalized Additive Models: An Introduction with R. New York: Chapman and Hall/CRC.

Yang Y, Hillebrand H, Lagisz M, Cleasby I, Nakagawa S. 2022. Low statistical power and overestimated anthropogenic impacts, exacerbated by publication bias, dominate field studies in global change biology. Global Change Biology 28, 969–989.

