## Supplementary Material for "HIGH FREQUENCY ROOT DYNAMICS: SAMPLING AND INTERPRETATION USING REPLICATED ROBOTIC MINIRHIZOTRONS"

### Supplementary Material for Online Publication

Supplementary Protocol S1: Full Instrument design

We built the RMR ourselves in our workshop. For imaging, the RMR uses a standard RGB digital camera. We changed this camera over the course of the experiments due to supply issues. In Experiment 1 we used an IDS 1005XS-C (IDS imaging Development Systems GmBH, Obersulm Germany). For Experiment 2 and 3, we used a DFK AFU050-L34 (TheImagingSource GmBH, Bremen Germany). There was a decrease in image quality in the second system. We used a customised fish-eye lens to allow the camera to focus with a short field of view (~ 3 cm) within the observatory. This had a slight impact on the distortion of field of view. We did not expect a meaningful structural distribution of roots on the same scale of individual images so we did not correct for this.

The mechanical and electrical components of the RMR were all purchased from suppliers but required some customisation for the precise use case. We used a LattePanda development board computer with integrated Arduino based coprocessor (ATmega32u4, LattePanda, Shanghai, China) and a WiFi antennae to allow remote access. The imaging unit was mounted on a custom-built chassis made of matt black plastic. We used two motors for 360° rotational (NEMA 17 stepper motor with custom 1:49 gear box) and a customized motorized linear rail system (RGS04K-M0394-A03, Haydon Kerk Motion Solutions, Waterbury CT, USA). This was controlled by the microcontroller alongside a 24 ~4500 k 'natural light' led ring (WS2812, Worldsemi, Dongguan, Guangdong, China) which was only switched on when images were taken and was angled away from the image subject to reduce reflectance. All exposed screw heads were painted black for the same purpose. Cable management was achieved by affixing cables to a rigid band only able to bend in the lateral direction. A power supply converter from 12V to 5V was used for the PC, camera and LED ring lights.

We set the timing of sampling through a manual timer switch (RS Pro time switch 1 channel, RS components, Corby, UK) operating the mechanical movement and image capture cycle through shell scripts which ran when the computer successfully booted up and when imaging was complete, shut down the computer. A full sampling cycle takes ~ 40 minutes, covering the entire tube and RMR takes 112 images per cycle, at 8 rotational and 14 lengthwise positions. Images were saved in .jpg formats onto removeable 128 GB SD cards; in order to obtain the images, the SD cards were swapped with blank alternatives during periods when the instruments were not powered. A log file was amended at the end of each imaging cycle which could be checked much faster than individual images and easily accessed over WiFi. A single image was less than 1 Mb, therefore the RMR could sample around 1100 cycles without the SD card being changed.

One sampling cycle from the RMR requires about 10 watt hours of power. Here in E1 and E2 and E4 the RMR ran on mains power, although on occasion was unplugged for several cycles due to other work in its greenhouse location, and/or minor modifications as in this development phase we still had concerns about cable management in the mechanical part of the device. These modifications did not affect positioning nor later image capture. In E3 the instruments ran on solar power which meant an occasional loss of data in rare extended bad weather when the battery did not recharge sufficiently. The images collected by the RMR overlap by 250 pixels in the lateral direction. For image processing, we trimmed the images to remove overlap so all pixels were independent.

This design of the RMR operates in a 1000 m long x 100 mm (96 mm internal) diameter observatory. This is comparable to the only other automatic minirhizotron systems used in published experiments (Iversen *et al.*, 2011; Svane *et al.*, 2019).

In the instrument revision between E3 and E4, we made three minor modifications to the RMR. First, we installed a GPS clock which could correct any drift in the BIOS clock due to temperatures (encountered in E3, described in the main manuscript). Second, we added a magnetic sensor to reduce the reliance on mechanical precision to reset the instrument between runs; this prevented a rare but fatal cable management issue if the instrument overshot its ‘resting’ position. Thirdly, we changed the camera to a IDS 1007XS-C (IDS imaging Development Systems GmBH, Obersulm Germany).

Supplementary Figure S1: Sub-daily data and root diameter artefact

With the amount of training we performed (and hence, possibly improvable with more training effort), we encountered an artefact in the sub-daily data. This occurred when there were roots present in the mesocosm, segmented root cover was lower in periods of daylight than darkness (contrast midday and midnight in figure S1).

When we examined the whole segmented dataset, we found that this effect was strongest on the sides of the mesocosm and almost non-existent on the top. We therefore think that this effect was unlikely to be due to 1) sensor degradation, which should not vary within a single sampling routine which passed through a sequence of rotational positions without sampling any one angle completely before any other, 2) light 'leaking' through the shallow soil surface, which should affect the top of the mesocosm more than the bottom nor 3) poor tolerance of the positioning system, where errors should not occur in a structured fashion. Because this effect also only began to occur at levels of higher root biomass, and was difficult to distinguish with the native eye (Figure S1), we hypothesise that this may be due to changes in the soil appearance linked to diurnal depletion of root zone water linked to transpiration or potentially diurnal variation in root diameter (Huck *et al.*, 1970).

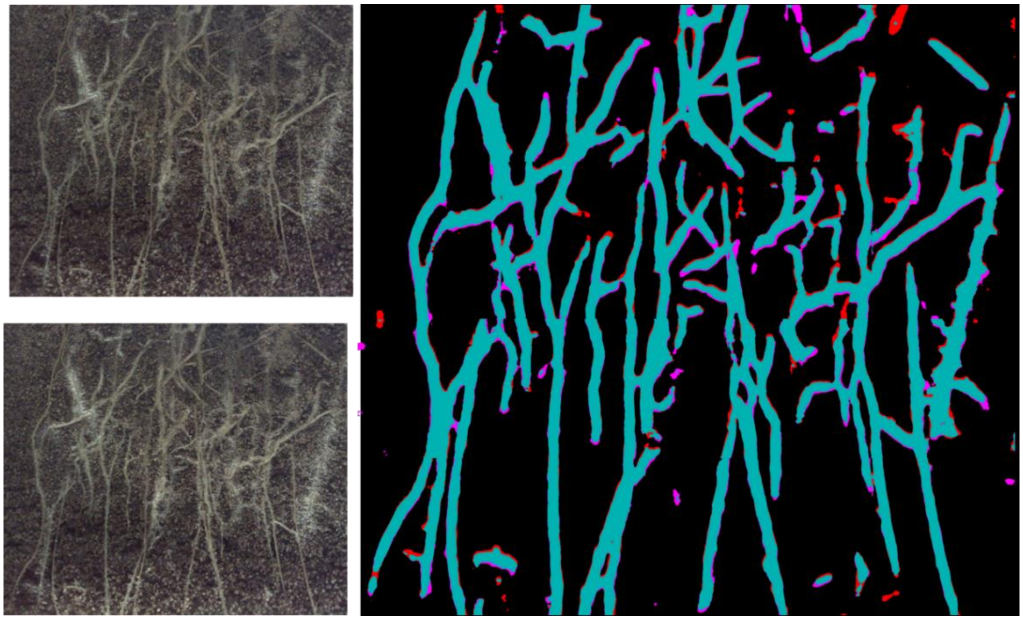

Figure S1 – Contrasting sequential day (from image top left) and night segmentations (from image bottom left). Areas in blue are root segmentations in common between a midday and midnight. Red is areas which were identified as roots at night, and soil during the day, while purple is areas that were identified as soil at night and roots during the day. Overall there are also more purple than red segmentations. This effect is common throughout the dataset.

<u>Supplementary Figure S2: Validation against Manual Measurements for Experiment 2, 3, 4.</u>

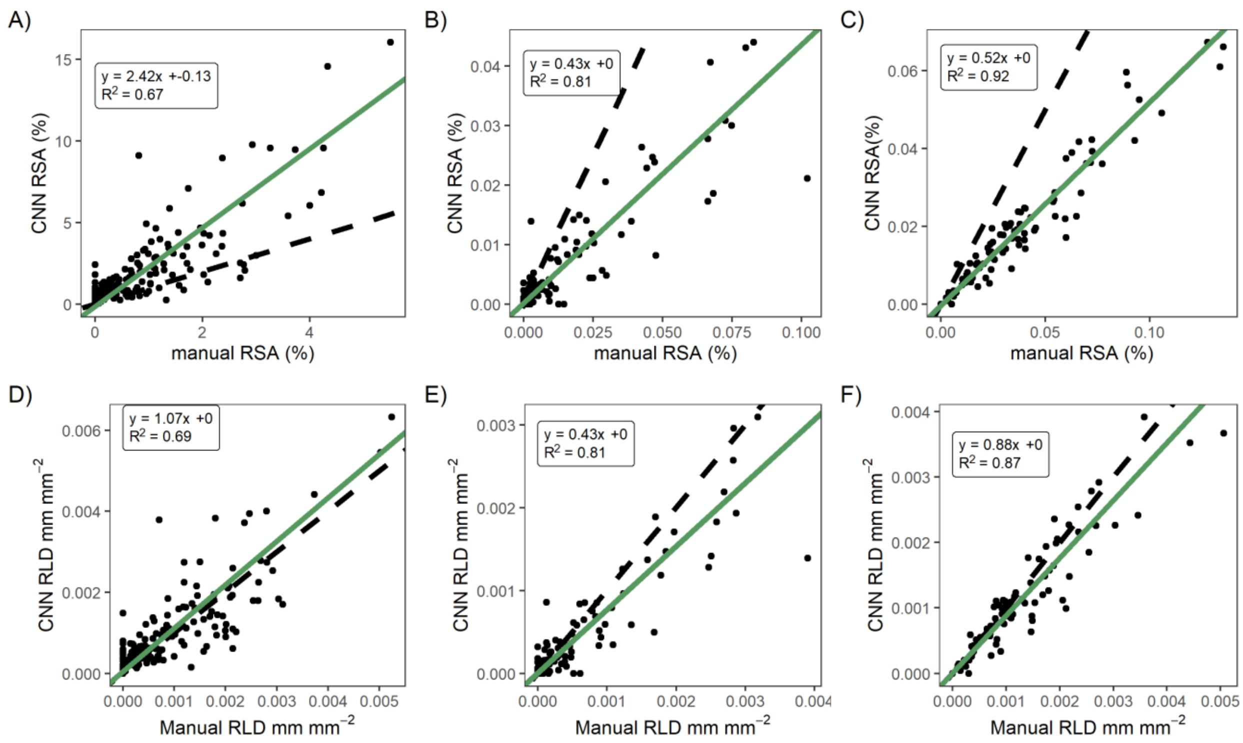


Figure S2: Image-level validation for Experiment 2, 3, and 4. The CNN consistently either over or under-estimated the manual RSA, which could be due to the model training or the validation data R we used. RLD was closer to 1:1.

#### Supplementary Figure S3: Consistency between mesocosm units

In experiment 2, we were able to segment a similar time series from all 8 mesocosms, with some variation as expected on each individual tube. Root cover was between 3 and 4 % of the images at their maximum and peaked around the same time, shortly after cessation of watering. Universally, root cover declined more slowly and later than GCC (Figure S5)

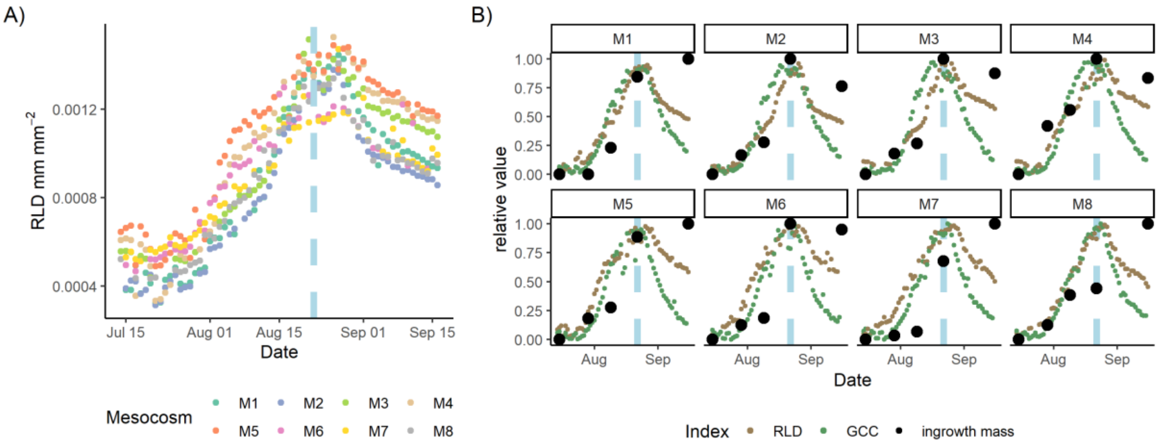


Figure S3: Comparison of absolute RSA from all mesocosm units in E2 (mesocosm names T101 through T108). Panel a) shows absolute mean RSA per mesocosm, absolute difference between mesocosms around 1 % of the total image or 25 % of the total area covered Panel b) shows comparison of GCC and RSA from all 8 mesocosm units. GCC matched RSA well in all cases and both indexes peaked after watering finished (blue line in both panels). In most mesocosms GCC reached its maximum RSA. Generally, the match between RSA and ingrowth root mass, and root cover and GCC was better at the start than the end of the experiment.


**Huck MG, Klepper B, Taylor HM**. 1970. Diurnal Variations in Root Diameter. Plant Physiology **45**, 529–530.

**Iversen CM, Murphy MT, Allen MF, Childs J, Eissenstat DM, a. Lilleskov E, Sarjala TM, Sloan VL, Sullivan PF**. 2011. Advancing the Use of Minirhizotrons in Wetlands. Plant and Soil **352**, 23–39.

**Svane SF, Dam EB, Carstensen JM, Thorup-Kristensen K**. 2019. A multispectral camera system for automated minirhizotron image analysis. Plant and Soil **441**, 657–672.
